# Human microphysiological model of dorsal root ganglion-spinal cord dorsal horn circuitry recapitulates opioid induced effects

**DOI:** 10.1101/2025.08.31.673344

**Authors:** Kevin J. Pollard, Frank R. Seipel, Nisha R. Iyer, Alex Bosak, Randolph S. Ashton, Michael J. Moore

## Abstract

Microphysiological systems (MPSs) are engineered, in vitro platforms which have been established as viable alternatives to animal models for pre-clinical research with unique advantages over conventional model systems. Many MPSs utilize 3-dimensional (3D) tissue constructs that enable biomimetic cell-cell interactions, allow for extended culture periods, and provide the time necessary for the emergence of physical and physiological characteristics of more mature tissues. Here, we present a novel MPS using human induced pluripotent stem cell (hiPSC)-derived spinal cord dorsal horn (SCDH) spheroids co-cultured with hiPSC-derived dorsal root ganglion (DRG) sensory spheroids in a microengineered hydrogel system to create a “connectoid” model of afferent pain circuitry. SCDH spheroids were functionally innervated by peripheral sensory neurons, and prolonged maturation of hiPSC-derived SCDH neurons within the connectoid system enabled derivation of crucial late-born cell types unattainable using 2D differentiations. Furthermore, hiPSC-derived SCDH spheroids spontaneously generate rhythmic, complex, synaptically-driven electrophysiological waveforms that are disinhibited by morphine exposure, consistent with spinal mechanisms of opioid-induced pruritus and hypersensitivity.

**One Sentence Summary:** hiPSC-derived afferent sensory circuitry model, with NK1R+ spinal cord dorsal horn neurons, yields electrophysiologically mimetic response to opioids.

## INTRODUCTION

Circuitry connecting the peripheral dorsal root ganglia (DRG) and the central spinal cord dorsal horn (SCDH) is a key junction in the propagation of sensory signals. This interface has been identified as a promising therapeutic target for both chronic pain and itch (*1, 2*), which are complex perceptions involving overlapping layers of signal processing throughout the nervous system. Normal signal propagation initiates with noxious stimulation of peripheral nociceptors (pain) or pruriceptors (itch), whose cell bodies reside in the dorsal root ganglion (DRG) (*3*). These signals are transmitted to the central nervous system (CNS) via synaptic transmission between DRG- sensory neurons and interneurons (INs) which populate the superficial laminae (I-III) of the spinal cord dorsal horn (SCDH) (*4, 5*). One of the key neurotransmitters involved in pain and itch signaling in the SCDH is substance P (SP), which is released by DRG-sensory neurons following noxious stimuli and bound by the neurokinin-1 receptor (NK1R), which is expressed on projecting SCDH-INs (*6, 7*). Upon NK1R activation, pain and itch signals can interact with each other through spinal interneuron circuitry (*8, 9*) and are integrated with concomitant, non-noxious afferent sensory signals as well as descending emotional and psychological feedback from the brain (*10–14*). The modulated sensory signals then project through the ventral spinal cord to local motor networks to mediate subconscious reflexes and ascend ventrolaterally where they are further modulated in the brain stem and thalamus before projecting to the cortex for conscious perception (*8, 9*).

The SCDH is normally maintained under inhibitory control by descending input and local Ins (*10*). Loss of this inhibitory tone is known to contribute to the development of chronic pain and itch (*11, 12, 15*). SP activation of NK1R strengthens synapses in the SCDH and sensitizes them, likely playing an important role in pain and itch chronicity (*16, 17*). Opioids remain a primary treatment for chronic pain and provide analgesia in part by activating µ-opioid receptors at the DRG-SCDH interface (*18*). However, reliance on addictive opioid-based therapeutics has contributed to over 700,000 opioid overdose deaths in the past two decades (*19*). Interestingly, NK1R antagonists have been widely explored as treatments for pruritus (*2, 7, 20*), and they could also be a viable target for novel non-opioid pain therapeutics (*6, 7*) due to well-documented crosstalk. For example, painful stimuli inhibit spinal itch circuits through synaptic activation of SCDH-INs and/or release of modulatory neuropeptides including the endogenous opioid dynorphin (*13, 14*). Moreover, increased itch sensation is a side-effect of opioid treatment, especially if applied neuraxially (*21*). However, while NK1R-targeting drugs to treat both pain and itch have shown pre-clinical promise, clinical trials for NK1R antagonists to treat itch have had mixed results (*20*), and NK1R-based therapeutics have not produced effective analgesia in humans (*17*). This lack of translational success brings into question the effectiveness of traditional pain and itch preclinical models.

Rodents are the primary preclinical model system used to evaluate novel analgesics and pruritus therapeutics but translation of their results to humans is confounded by differences in patterns of innervation within SCDH circuits, expression of pain and itch-related transcripts, and developmental trajectory of neurons and glia (*22–25*) reviewed in (*26*). While significant progress has been made in the generation of DRG-sensory neurons from human pluripotent stem cells (hPSCs) (*27, 28*), direct modeling of relevant DRG-SCDH circuitry has been very limited. In fact, only one recent publication has demonstrated derivation of NK1R+ SCDH-INs in a four-organoid assembloid system to model the ascending neural sensory pathways (*29*). While impressive in its comprehensive nature, the scalability of this system as a preclinical model for drug discovery may be limited.

Here, we describe a scalable microphysiological system (MPS) of lower afferent DRG-SCDH signaling generated directly from cryopreserved, hPSC-derived cell banks in a microengineered hydrogel device to create a connectoid between DRG and SCDH spheroids. By Day 35 of MPS culture, single cell RNA sequencing revealed that the model’s cellular composition can generate the SCDH and DRG cellular phenotypes required for recapitulating afferent pain and itch synaptic circuitry. Upon further maturation in long-term co-culture (> ∼Day 63 of MPS culture), late-born, NK1R+, laminae I/II neurons emerged in the SCDH spheroid, and this was concurrent with emergence of spontaneous, complex patterns of synaptic burst firing within the SCDH spheroid themselves, which required glutamatergic neurotransmission and were independent of DRG innervation. Furthermore, we observed that acute morphine exposure predictably disinhibited NK1R+ glutamatergic SCDH burst firing, mimicking acute bicuculline-mediate inhibition of GABAergic INs. Thus, our hPSC-derived DRG-SCDH MPS models morphine’s GABAergic-mediated disinhibition of SCDH circuitry observed *in vivo* (*30*). Loss of GABAergic inhibition, and resulting spinal cord circuit hypersensitivity, has been implicated in conditions of excessive sensation including both opioid-induced pruritus (*31*) and paradoxical pain hypersensitization (*32*). These results indicate that this DRG-SCDH connectoid model could facilitate discovery of novel chronic pain and itch therapeutics.

## RESULTS

### Derivation of spinal cord dorsal horn (SCDH) neuronal cultures from hPSCs

Human iPSC-derived DRG sensory neurons can be acquired commercially and have been shown to contain neurons with both nociceptive and pruriceptive characteristics (*28, 33*) (Fig. S1). Thus, we sought to generate a SCDH population from hiPSCs (iMR90-4), which could contain the requisite excitatory and inhibitory interneurons (INs) responsible for processing and relaying sensory signals into the central nervous system (*8*) (Fig. 1A). During development, these cells emerge from dorso-intermediate progenitors in the neural tube (pd4/pdL/pd5), which are patterned by TGF-β-independent signaling pathways and are defined by cross-repressive transcriptional interactions (Fig. 1B) (*9*). Neuronal birthdate further distinguishes different SCDH-INs. Early-born neurons (dI4/dI5) express Onecut2 and Zfhx3, and late-born neurons (dILa/dILb) express Nfia and NeuroD2 (*34*). These late-born neurons ultimately migrate from intermediate spinal progenitors to populate the superficial dorsal horn (laminae I-III) where they receive, modulate, and further transmit pain and itch signals from the DRG (Fig. 1A,B) (*34, 35*).

**Fig. 1.**
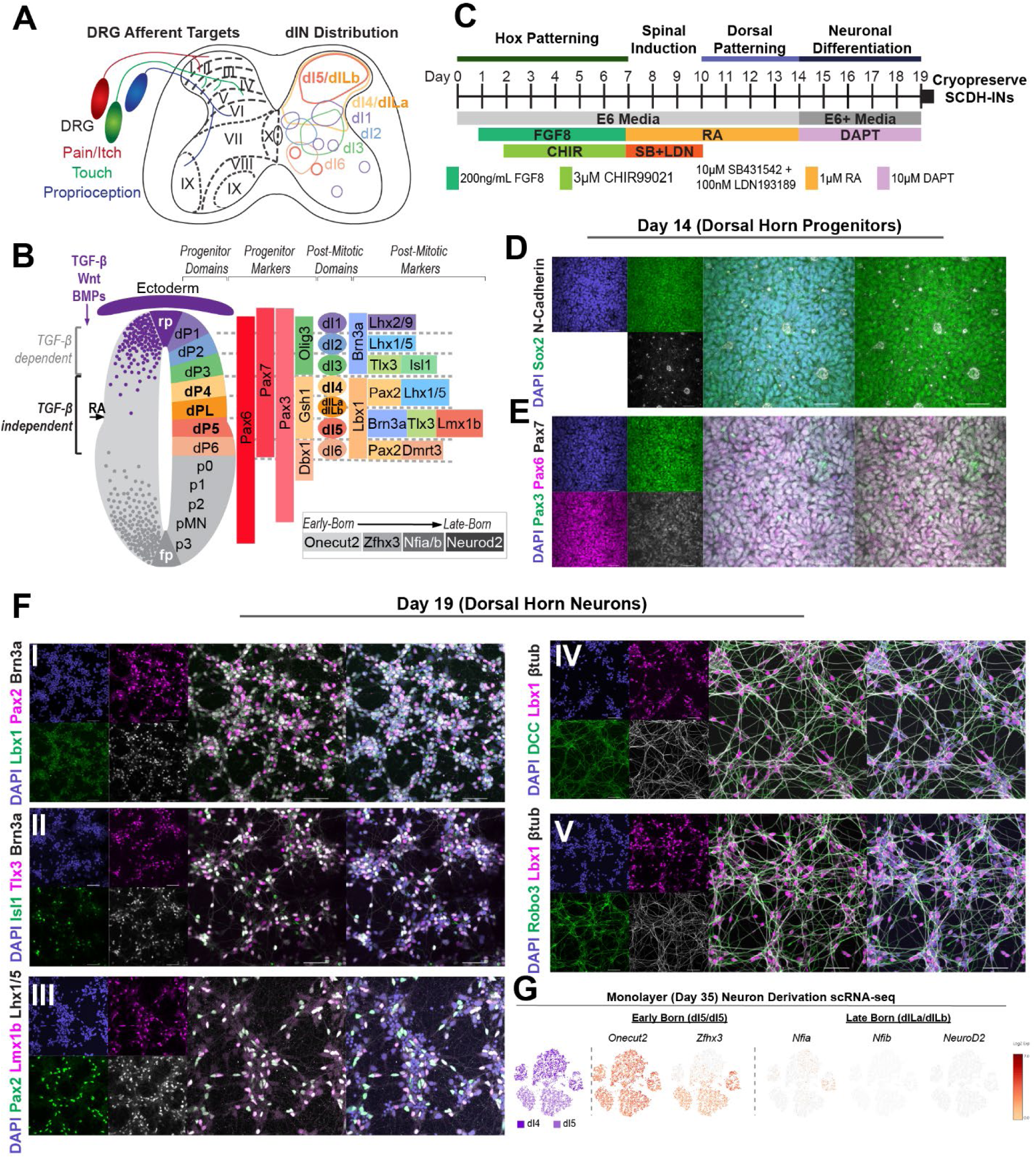
iPSC Derivation of Human Spinal Cord Dorsal Horn Neurons: **A)** Schematic demonstrating innervation targets of DRG sensory neurons in the SCDH. **B)** Developmental origin of SCDH neurons. **C)** Differentiation scheme for Dorsal Horn Progenitors (halt differentiation on day 14) and SCDH-INs (cryopreserve on day 19). **D-E)** Immunohistochemical staining (IHC) of day 14 Dorsal Horn Progenitors indicating **(D)** neural progenitor capacity (Sox2), the formation of polarized rosettes (N-Cadherin), and **(E)** dorsal spinal progenitor identity (Pax6, Pax7, Pax3). **F)** IHC of day 19 SCDH neurons. **(i-iii)** Staining for subtype markers indicating Lbx1+ SCDH-INs with both inhibitory (Pax2+, Lhx1/5+) and excitatory (Brn3a+,Tlx3+,Lmx1b+) populations and low prevalence of dI3 neurons (Lbx1-, Tlx3+, Isl1+) (see Fig. 1B). **(iv,v)** Staining for neuronal identity (β-tubulin) and markers of axonal guidance (Robo3, DCC). **G)** scRNA-seq analysis of monolayer-derived SCDH-INs (*36*). dI4/5 neurons broadly express the early-born marker Onecut2 with dI5 neurons additionally expressing Zfhx3, another marker of early birthdate in CNS neurons. Late-born markers Nfia, Nfib, and NeuroD2 are not broadly expressed in monolayer derived SCDH-INs.

To derive a transcriptionally appropriate population of human SCDH-INs, we adapted our previously-published 2D-monolayer protocol for differentiating hPSCs into a diverse spectrum of regionally distinct spinal neuron cultures with discrete *HOX* profiles (*36*) (Fig. 1C). By patterning lower cervical spinal progenitors with only retinoic acid (RA), cultures were directed to intermediate neural tube fates. By day 14, SCDH progenitors had formed N-cadherin+ neural rosettes (Fig. 1D) and were PAX6+/PAX3+/PAX7+ (Fig. 1E). Rapid neuronal differentiation using DAPT treatment resulted in LBX1+ SCDH-INs that were distributed between excitatory (dI5/dILb; BRN3A+/TLX3+/LMX1B+) and inhibitory (dI4/dILa; PAX2+/LHX1/5+) cell types (Fig. 1F). Moreover, SCDH-INs expressed key functional markers of axon guidance, DCC (*37*) and ROBO3 (*38*), that are characteristic of commissural projection neurons that cross the ventral midline (*9, 39*). ROBO3 is specifically restricted to dorso-intermediate neurons (dI4-dI6), which suggests that our SCDH-INs appropriately express both transcriptional and functional markers particular to this population. While our monolayer differentiation scheme enables the generation of excitatory and inhibitory SCDH-IN populations, the resulting dI4/dI5 neurons do not express the late-born transcription factors which are characteristic of the dILa/dILb population that ultimately populates the superficial SCDH (Fig. 1G) (*36*).

### Generation of a human microphysiological model of SCDH Innervation by DRG peripheral sensory neurons

As recently observed in neural assembloid culture (*29*), we hypothesized that maintaining our hPSC-derived dorso-intermediate neuronal cultures in long term-3D culture could extend maintenance of the progenitor pool and could enable generation of a late-born SCDH-IN population. Additionally, we sought to recapitulate afferent DRG-SCDH circuitry by performing this 3D culture in an MPS alongside spheroids generated from hPSC-derived DRG sensory neurons. Thus, spheroids generated from cryopreserved day-19 SCDH-IN cultures (Fig. 1C) and commercially available DRG sensory neurons (Anatomic, Inc.) were seeded into a growth-restrictive PEG hydrogel mold which was then filled with growth-permissive Matrigel^®^ (Fig. 2A). We previously showed that this MPS design could permit long-term tissue growth, cellular maturation, and directional innervation of embryonic rat-derived DRG and SCDH spheroids (*40*). Over time, the gross tissue-level properties of a lower afferent pain circuit emerged (Fig. 2B,C) in the “connectoid” system. Somata of mature SCDH neurons and DRG sensory neurons showed clear morphological differences at 49 Days in gel (DIG) (Fig. 2Ci,ii). DRG somata are large with a single axon extending distally while SCDH somata are small and extend extensive, branching, neuritic arbors. Both tissue types express the presynaptic protein Synapsin-I but SCDH neurons expresses more of the post-synaptic protein PSD-95 (Fig. 2Ciii,iv). This qualitatively suggests that functional pre-to-post synaptic connections are preferentially occurring in the SCDH. The GABAergic marker, VGAT, is preferentially expressed in the SCDH spheroid while the glutamatergic marker, Vglut, is similarly expressed by both DRG and SCDH neurons consistent with the purely glutamatergic nature of Peripherin+ DRG sensory neurons and mixed presence of glutamatergic and GABAergic interneurons in the SCDH-IN *vivo* (*34*) (Fig. 2Cv,vi, vii). Also, GFAP+ glial cells spontaneously emerge after prolonged MPS culture and populate both the SCDH and DRG spheroids (Fig. 2Cviii).

**Fig. 2.**
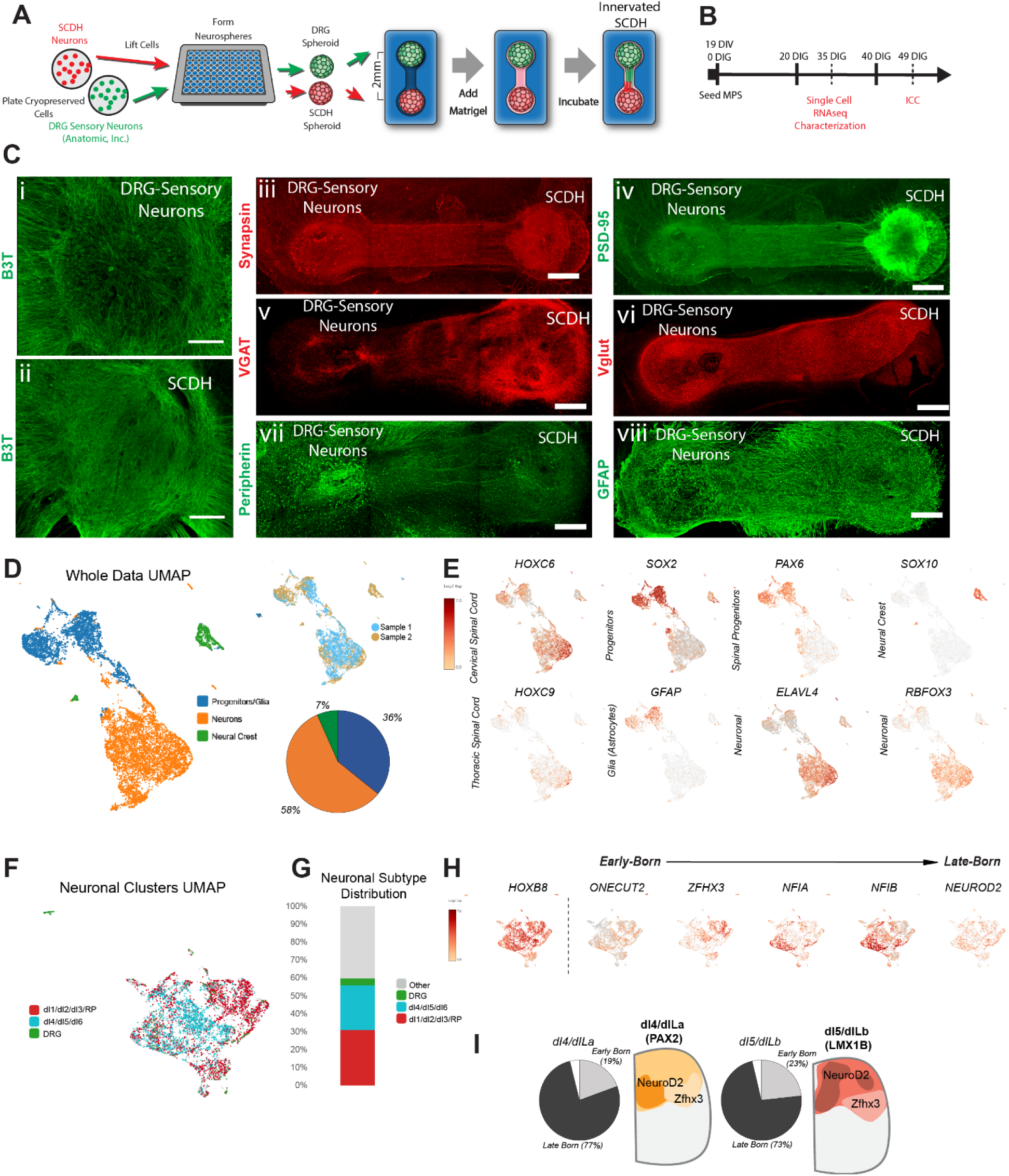
Characterization of DRG-SCDH MPS: **A)** Schematic demonstrating culture and generation of DRG-SCDH MPS. **B)** Timeline for MPS culture and analysis. **C)** IHC demonstrating the gross morphological properties of the DRG-SCDH MPS. Stains shown include ***(i,ii)*** pan-neuronal marker, β-III-tubulin (B3T), ***(iii)*** presynaptic marker, synapsin (syn), ***(iv)*** postsynaptic marker, post-synaptic density protein-95 (PSD-95), ***(v)*** GABAergic cell marker vesicular GABA transporter (VGAT), ***(vi)*** glutamatergic cell marker, vesicular glutamate transporter 2 (VGLUT1), ***(vii)*** peripheral neuron marker, peripherin, and ***(viii)*** glial cell marker glial fibrillary acid protein (GFAP). Scale bars = 200 µm. **(D-I)** scRNA-Seq analysis of SCDH-INs at 35 DIG in MPS culture. **(D,E)** UMAP projection of complete UMI-Filtered dataset indicating a split population of neurons and progenitors/glia with a small prevalence of neural crest cells. **(F,G)** Re-clustering of the neuronal cells to reveal the distribution of neuronal subtypes at 35 DIG. **(H)** Neurons from the MPS at 5 weeks express upper-laminae sensory marker HoxB8 and have widespread expression of late-born markers Nfia, Nfib, and NeuroD2 which are not observed in monolayer only derivations (Fig. 1). **(I)** Distribution of early and late born cells making up MPS-derived inhibitory (dI4/dILa) and excitatory (dI5/dILb) SCDH-IN populations at 35 DIG and schematics demonstrating their spatial distribution in the in vivo SCDH.

### Single Cell RNA-seq Characterization of DRG-SCDH MPS and comparison to 2D monolayer derivation

To characterize the cellular composition of our DRG-SCDH MPS, single cell RNA sequencing (scRNA-seq) of 11,321 cells from 35 DIG cocultures was conducted (Fig. 2B). This analysis showed that the spheroids differentiate exclusively into neurons (ELAVL4+/RBFOX3+), neural progenitors (SOX2+/PAX6+), glia (GFAP+), and neural-crest (SOX10+) progeny corresponding to the cervicothoracic spinal cord (HOXC6+/HOXC9+) and DRG, without contamination of extra-nervous system cell types (Fig. 2D,E). Isolation and re-clustering of neuronal cells reveals enrichment of both excitatory and inhibitory DRG and SCDH-INs (Fig. 2F,G, Fig. S1A,B). Importantly, SCDH cells express HOXB8, a critical marker for upper laminae sensory neuron development (*41*), and express both early-born (ONECUT2, ZFHX3) and late-born markers (NFIB, NEUROD2) (Fig. 2H). The dI4/dILa and dI5/dILb populations can be further stratified by their expression of these temporal markers, which allows for extrapolation of laminar organization (*34*) (Fig. 2I). Previously published scRNA-seq data comprising 2D monolayer, hPSC-derived, dI4 and dI5 INs failed to express late-born transcription factors (1G) (*36*), which suggests that extended 3D culture is necessary for the development of a transcriptionally- appropriate late-born superficial SCDH population.

We hypothesize that the difference in late born marker expression between our previously derived 2D monolayer (*36*) and current 3D MPS SCDH cultures is due to the maintenance of a Sox2+/Pax6+ progenitor pool (Fig. 2D,E). Thus, we further compared the scRNA-seq profile of neural cells generated in the 3D-MPS culture versus using our previous 2D monolayer differentiation (*36*) (Fig. 3A). We observed that the MPS and 2D differentiated cells segregate into distinct populations (Fig. 3B). Next, neurotransmitter-positive neurons were isolated and re-clustered independently producing 9 distinct MPS culture and 21 distinct 2D-culture neuronal subtypes. Also, the clusters were categorized by expression of distinct neurotransmitter systems and by regionally-specific gene targets using a standard dotplot in R/Seurat (Fig. 3C). Clusters recovered from the MPS were characterized as dorsal glutamatergic interneurons (MPS-dorsal- Glut), dorsal GABAergic neurons (MPS-dorsal-GABA), or cells that maintain a progenitor phenotype (MPS-prog), with no evidence of the ventral interneuron, dopaminergic, or cholinergic cell types purposefully generated using Shh+ patterning in the prior 2D differentiation protocol (*36*). MPS-prog1-3 clusters expressed high levels progenitor markers GFAP, SOX2, PAX6, and SOX10 and relatively reduced expression of mature neuronal, interneuron, and neurotransmitter system markers. This is consistent with maintenance of a progenitor pool in the 3D-MPS culture, and it is absent in our prior 2D monolayer neuronal cultures (*36*) (Fig. 3C). Also, the MPS-matured neuronal clusters expressed higher levels of cervical and thoracic spinal cord markers HOXC6 and HOXC9 and a more complete subset of dI1-6 identity markers including the dI2-specific marker FOXD3 (MPS-Glut1 and MPS-Glut2), the dI4/6 marker PAX8 (MPS-GABA1), and the dI6-specific marker BHLHE22 (MPS- GABA2), none of which were expressed to a significant degree in the analogous 2D monolayer clusters (Fig. 3C). This suggests that prolonged maturation in 3D-MPS culture produces a more diverse neuronal population than our directed monolayer differentiation method, perhaps better recapitulating *in vivo* developing spinal cord and enabling the generation of late-born SCDH neuronal subtypes (Fig. 2G).

**Fig. 3.**
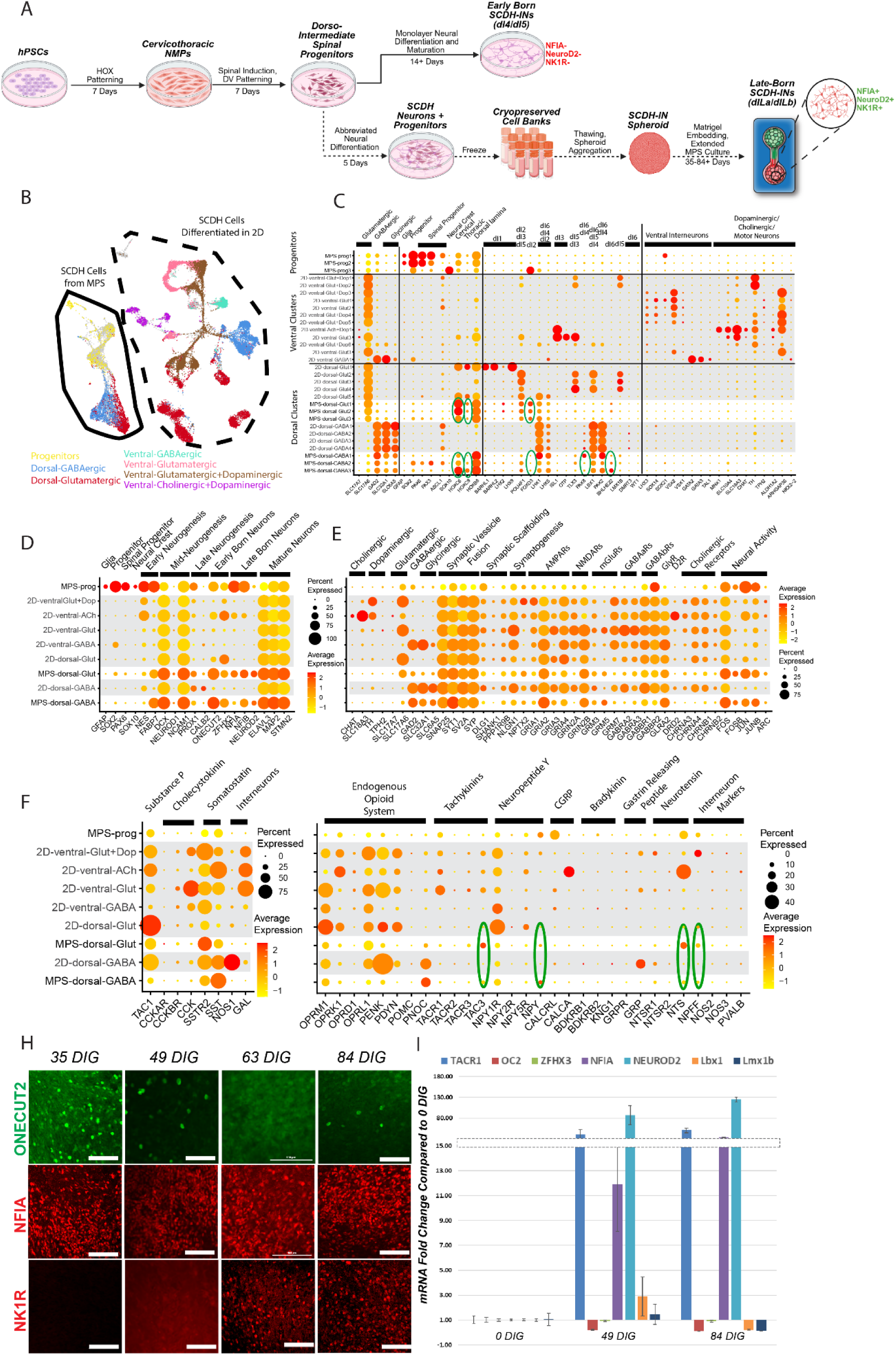
Comparative analysis of gene expression patterns in 2D and MPS-matured SCDH-Interneurons. **(A)** Schematic of differentiation timelines for 2D and MPS-matured SCDH-IN populations. **(B)** 2D and MPS-derived SCDH-Interneuron single cell sequencing datasets were combined and re-clustered to define distinct neuronal populations. **(C)** 2D differentiated ventral spinal cord neurons clustered into glutamatergic, GABAergic, dopaminergic, and cholinergic populations. 2D differentiated dorsal spinal cord interneurons clustered into glutamatergic and GABAergic populations. MPS-derived SCDH-IN populations clustered into glutamatergic and GABAergic interneuron populations as well as three progenitor populations. **(D)** Fig. 3C clusters were collapsed based on dorsal/ventral identity and expression of distinct neurotransmitter systems. **(E)** Expression of proteins necessary for functional neurotransmission including markers of neurotransmitter systems, synaptic vesicle fusion, synaptic scaffolding, and synaptogenesis. **(F)** Expression of pain related neuromodulators and neuromodulator receptors. **(H)** Immunofluorescent staining of spheroids at 35, 49, 63, 84 DIG for temporal markers and the substance P receptor (NKR1). Scale bars=100µm. **(I)** qRT-PCR analysis of Week 7 and 12 spheroids for dI4/5 subtype markers (LBX1, LMX1B), early-born markers (*ONECUT2, ZFHX3*), late-born markers (*NFIA, NEUROD2*), and the substance P receptor (*TACR1*). Panel **(A)** Created in BioRender. https://BioRender.com/q032tb6

### Expression of mature circuitry and neuromodulator markers implicated in pain and itch perception

We further compared the expression of genes relevant to the function of synaptically-connected dorsal spinal cord neural circuits. Clusters were collapsed according to neurotransmitter system, dorsal-ventral identity, and expression of genes related to neurogenesis, synaptic transmission, and pain and itch-related neuromodulators (Fig. 3D-G). MPS progenitors expressed the highest levels of early neurogenesis genes, while MPS glutamatergic and GABAergic interneurons expressed the highest levels of mid-to-late neurogenesis, early-born neuron, late-born neuron, and mature neuron markers (Fig. 3D). Both SCDH neuronal populations derived in 2D or 3D-MPS culture express the basic functional proteins required for synaptic transmission including genes necessary for synaptic vesicle formation, synaptic vesicle fusion, synaptic scaffolding, and synaptogenesis (Fig. 3E). Moreover, MPS neurons expressed higher levels of genes related to recent neuronal activity (FOS, FOSB, JUN, JUNB, and ARC) suggesting they may be more active at baseline than those matured in our prior 2D culture alone.

The superficial SCDH is heavily populated by glutamatergic and GABAergic interneurons, and GABAergic interneurons are also commonly glycinergic (*4, 5*). Here we found that both 2D and MPS-matured SCDH-Interneuron populations were also largely glutamatergic (expressing *SLC17A6*) or GABAergic (expressing *GAD2*), and GABAergic clusters were also glycinergic (expressing *SLC32A1* and *SLC6A5*) (Fig 3E). Glutamate, GABA, and glycine receptors were also expressed by both 2D and MPS-matured SCDH-INs with some differences in receptor subtype-specific expression. MPS neurons expressed a higher *GRIN2B* to *GRIN2A* ratio relative to 2D dorsal clusters, which is characteristic of active synaptogenesis (*42, 43*). MPS neurons expressed more *GRM5* (phospholipase C-activating) but less *GRM7* (cyclic AMP-inhibiting) likely altering the balance of second messenger signaling cascades after glutamate binding (*44*). Finally, MPS neurons expressed more *GABRA2* relative to *GABRA3*. GABA_A_ receptor subunit composition is known to affect ligand affinity and kinetics of activation and has been shown to alter the efficacy of analgesics that act through allosteric modulation of GABA receptors (*45*). In contrast, dopaminergic and cholinergic neuronal populations were largely absent from both 2D and MPS-matured SCDH-INs and dopamine and acetylcholine receptors were minimally expressed (Fig. 3E). *In vivo,* cholinergic and dopaminergic receptors are expressed and functional in the SCDH (*46, 47*) but these receptors are activated through long-distance projection neurons whose soma reside outside of the SCDH (*48, 49*).

Lastly, using scRNA-seq analysis, we characterized expression of pain and itch-related neuronal subtype markers present within 35 DIG MPS cultures (Fig. 3F,G). Consistent with descriptions in the rodent spinal cord dorsal horn (*30*), the mu opioid receptor (*OPRM1*) was expressed similarly by both MPS GABAergic and glutamatergic spinal cord interneurons. The kappa opioid receptor (*OPRK1*) was modestly expressed while the delta opioid receptor (*OPRD1*) was minimally detectable in MPS interneurons. Expression of the endogenous opioid gene PENK was reduced but detectable in MPS interneurons relative to analogous 2D clusters. Endogenous opioid genes *PDYN* and *POMC* were minimally detectable. Expression of the opioid-related ligand, nociceptin *(PNOC*) was highly expressed by 2D and MPS-derived dorsal GABAergic neurons while its receptor, nociception opioid peptide receptor (NOR/*OPRL1*) was expressed in their dorsal glutamatergic neuron counterparts. An array of neuromodulators and neuropeptides have also been implicated in the processing of pain and itch in the SCDH (*8*). Here we found that the neuromodulators substance P, cholecystokinin (*CCK*), calcitonin gene-related peptide (CGRP/*CALCA*), bradykinin, and gastrin releasing peptide were minimally expressed in both 2D and MPS-matured SCDH-INs while the neuromodulators neurokinin B (*TAC3*), neurotensin (*NTS*), and neuropeptide FF (*NPFF*) all showed increased expression in MPS-derived INs (Fig. 3G). The cognate receptors for each of these neuromodulators were lowly expressed in both 2D and MPS-derived SCDH-IN populations. These results suggest that although MPS-based differentiation produces a more diverse population of dorsal spinal interneurons at 35 DIG, expression of the neuromodulator systems involved in regulation of these spinal circuits remains incomplete.

### Expression of pain- and itch-specific TACR1+ neurons emerge with extended MPS culture

Expression of *TACR1*, the gene coding for the Substance P receptor NK1R, was largely absent in this 35 DIG SCDH neuronal population in 3D-MPS culture (Fig. 3H, Fig. S1A). We hypothesized that further maturation of SCDH spheroids within the 3D MPS could allow for the emergence of an NK1R+ SCDH population. Immunostaining confirmed the presence of late-born marker NFIA in 35 DIG SCDH spheroids and its maintenance through 84 DIG (Fig 3H). Early-born marker ONECUT2 expression decreases at later time-points. Remarkably, it took until 63+ DIG before widespread NK1R (*TACR1)* expression is observed and thereby validating our hypothesis (Fig 3H). qPCR analysis of 0, 49, and 84 DIG cultures aligned with our immunostaining analysis, with elevated levels of late-born transcription factors (*NFIA*, *NEUROD2*) and *TACR1* at 49 and 84 DIG (Fig. 3I). Additionally, SCDH neurons maintained protein-level expression of dorsal horn subtype markers Lbx1 and Brn3a throughout the culture time course (Fig. S1D-G).

### Early 3D MPS maturation: SCDH synaptic circuitry evoked by glutamatergic excitation and disinhibited by morphine

With the 3D MPS’s DRG-SCDH mimetic cellular composition confirmed, we proceeded to characterize the system’s electrophysiological behavior. Functional physical connections between the peripheral and spinal spheroids form after about 40 DIG. Between 40 – 70 DIG, electrical stimulation of innervating DRG spheroid produces a complex and prolonged electrical response within SCDH spheroid (Fig. 4A,B). This synaptically-evoked waveform is comprised of a fast, negative-going component followed by one or more relatively prolonged, positive-going deflections (Fig. 4B, Sham). Similar to our previous description using an analogous embryonic rat-derived DRG-SCDH MPS (*40*), application of excessive concentrations of morphine (∼1 mM) has an inhibitory effect (Fig. S2). However, application of the lowest effective dose of morphine (100 μM) selectively enhances the late portion of the evoked P-wave (Fig. 4B,C). This is mimicked by application of the ionotropic GABA_A_ receptor antagonist, bicuculline (BCC) (Fig. 4D,E), and abolished by application of the glutamatergic AMPA/kainate receptor antagonist, 6-cyano-7-nitroquinoxaline-2,3-dione (CNQX) (Fig. 4F,G). Consistent with the embryonic rat-derived MPS (*40*), these results indicate that activation of microphysiological SCDH circuitry requires glutamatergic neurotransmission, is limited by GABAergic neurotransmission, and is disinhibited by opioid receptor activation.

**Fig. 4.**
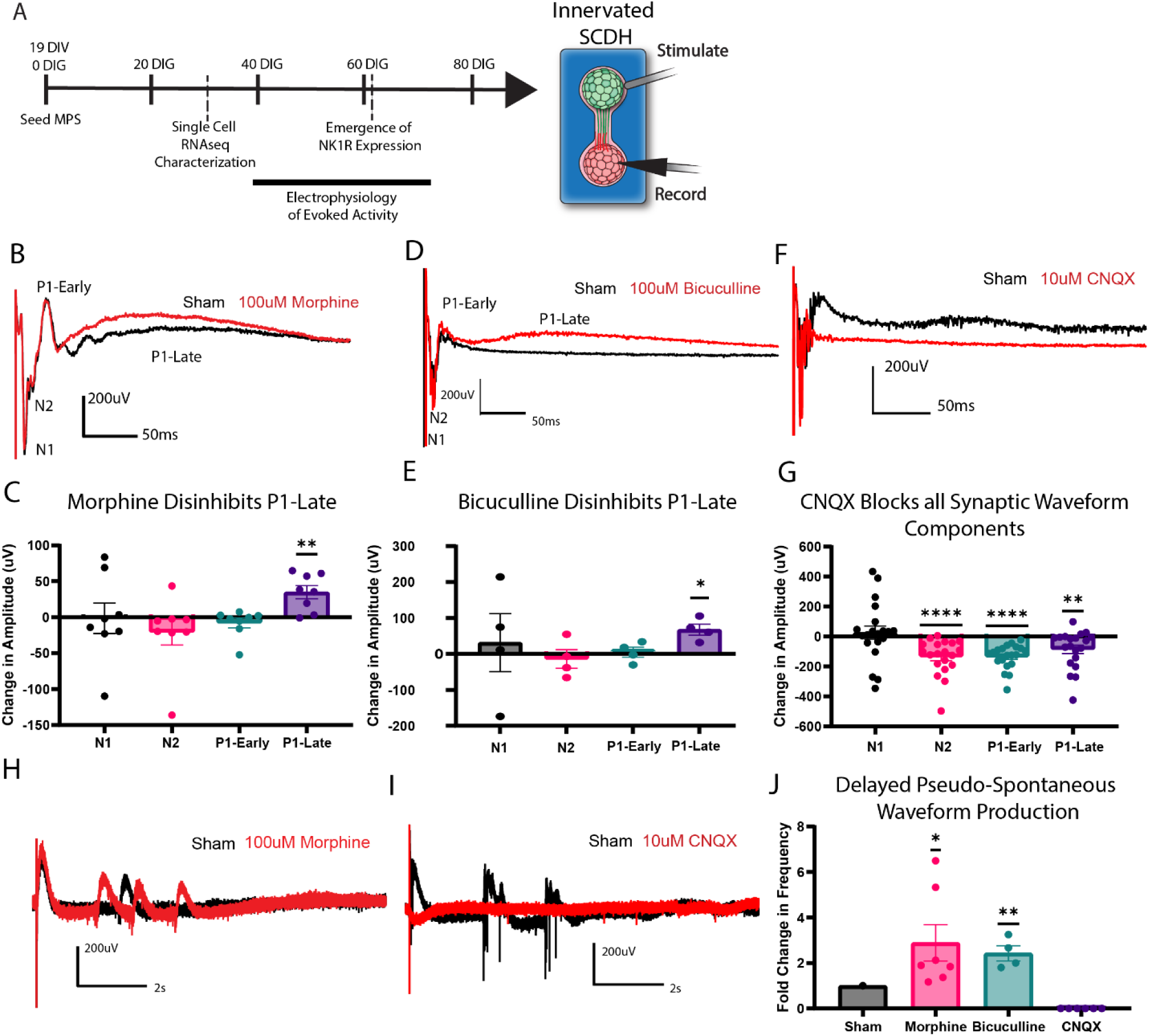
Synaptically-evoked SCDH activity is disinhibited by morphine. **(A)** Evoked activation of SCDH spheroid by electrical stimulation of innervating DRG spheroid was evaluated between 40 and 70 days in gel (DIG). **(B,C)** The late phase of P1 is selectively enhanced after application of 100µM morphine. **(D,E)** Direct inhibition of GABAergic synaptic transmission with bicuculline enhances P1-late potentials similar to morphine. **(F,G)** The N2, P1-early and -late components of the complex synaptic waveform are eliminated after blockade of AMPAR-dependent synaptic transmission with CNQX. **(H)** Pseudo-spontaneous positive-going field potentials continue to be produced in the SCDH several seconds after electrical stimulation and the frequency of spontaneous field potentials is increased following application of morphine and **(I)** eliminated following application of CNQX. **(J)** Quantification of pseudo-spontaneous complex waveform frequency in sham and morphine, bicuculline inhibition of GABAergic synaptic transmission, and CNQX AMPAR inhibition conditions. *p<0.05, **p<0.01, ****p<0.0001 per a one sample t-test vs. 0 (C,E,G) or 1 (J).

Additionally, a single DRG spheroid stimulation was often observed to evoke the production of multiple, repeated complex waveforms in the SCDH spheroid several seconds after the exogenous stimulation (Fig. 4H,I). The delayed production of these pseudo-spontaneous complex waveforms was also disinhibited by application of morphine and bicuculline and abolished by CNQX (Fig. 4H-J). This again suggests that opioid receptor activation increases the excitability of the microphysiological SCDH synaptic circuits similar to direct blockade of inhibitory GABAergic neurotransmission.

### Late 3D MPS maturation: SCDH synaptic circuits are rhythmically and spontaneously active and disinhibited by morphine

Corresponding to the emergence of NK1R+ SCDH neurons (Fig. 3H), consistent, concerted, and complex spontaneous waveform production emerges after ∼70 DIG, and the frequency of this spontaneous activity continued to steadily increase over time (Fig. 5A-C). Surprisingly, the presence of innervating DRG sensory neurons did not appear to alter the developmental trajectory of spontaneous activity (Fig. 5C). Like the synaptically-evoked waveforms, these spontaneous complex waveforms were relatively large in amplitude and resulted in prolonged positive deflections of the field potential beyond the timeframe of discrete action potentials or simple monosynaptic post-synaptic potentials (Fig. 5E,F). In contrast, spontaneous activity emerged at >100 DIG in DRG sensory MPS (Fig. 5B,D). Initial waveforms in the DRG spheroids are smaller in amplitude, were negative or biphasic field potentials, and only persisted for milliseconds, consistent with individual, discrete spiking behavior (Fig. 5D,G). Larger concerted waveforms did not appear until much later in the DRG spheroid (i.e., >200 DIG), and likely represented formation of artifactual synaptic connections due to prolonged MPS cultures.

**Fig. 5.**
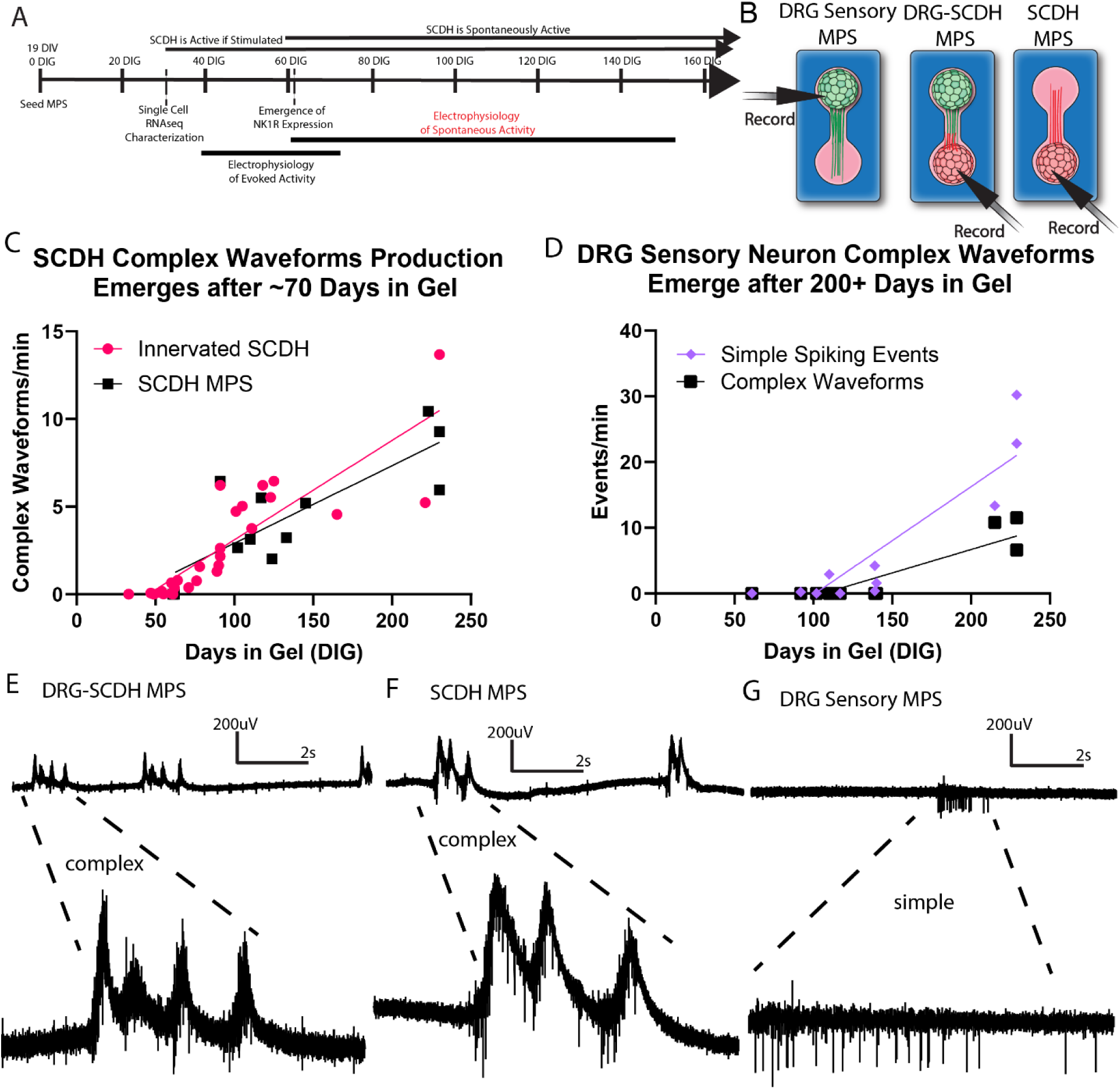
Spontaneous SCDH activity emerges after ∼70 DIG. (A) Time-course of evolution of spontaneous electrical activity in SCDH and DRG microphysiological models. (B) Recording electrodes are placed in nerve tissue and passively record spontaneous activity produced by the tissue in absence of any exogenous stimulation. (C) In SCDH spheroids, whether or not its innervated by a DRG spheroid, complex waveforms production begins after about ∼70 DIG and continues to increase in frequency up to 230 DIG. (D) Simple burst firing behavior was observed in DRG spheroid MPSs as early as 111 DIG, and complex waveform production was not observed until at least 200 DIG. (E,F) Complex burst in DRG-SCDH MPS or SCDH-MPS result in prolonged elevation of the extracellular field potential and occur as single or repetitive positive-going peaks separated by seconds or tens of seconds. (G) In contrast, the DRG MPS bursts are separated by tens of seconds but lack prolonged elevation of the extracellular field potential, are smaller in magnitude, and are either symmetrical about zero or primarily negative-going.

### Morphine effects on the SCDH spheroid’s glutamatergic spontaneous activity

Spontaneous SCDH activity was evaluated between 70 and 100 DIG (Fig. 6A) when spontaneous complex waveforms production was consistently identifiable (Fib. 5C). A paired recording paradigm was employed to evaluate spontaneous activity in the presence of different pharmacological compounds. During the recording period, one MPS was bathed in drug (morphine, CNQX, TTX, or ZD7288) while the paired control was bathed in vehicle artificial cerebrospinal fluid (ACSF) alone. Bioelectric field potential production was then passively recorded for one hour without any applied exogenous stimulation (Fig. 6B) and the frequency of waveforms produced was calculated. Morphine significantly increased the rate of spontaneous complex field potential production in SCDH spheroids of both DRG-SCDH MPS and SCDH MPS cultures aged 90-120 DIG (Fig. 6C,D,G). Furthermore, spontaneous activity was abolished after blockade of glutamatergic neurotransmission with CNQX (Fig. 6E,H) and by a 90-minute pretreatment with the HCN channel inhibitor, ZD7288, suggesting that these pacemaker-type channels are also necessary for periodic excitation of these synaptic circuits (Fig. 6I) similar to previous descriptions in *ex vivo* rodent SCDH tissues (*50, 51*). Simple spiking remained in the presence of CNQX that was abolished by further application of the voltage-gated sodium channel inhibitor, tetrodotoxin (TTX) (Fig. 6E,F). Similar CNQX-sensitive complex waveform production was also observed in an analogous MPS model of SCDH-Innervation comprised of embryonic rat DRG and SCDH spheroids (Fig. 6J,K). We did not observe analogous morphine- dependent potentiation of simple spiking or complex waveform production in DRG spheroids (Fig. S3). Together these results suggest that: spontaneous patterns of activity are generated within the SCDH spheroids themselves; spontaneous patterns are independent of interaction with innervating DRG sensory neurons; spontaneous events require HCN pacemaker channel activity; the resulting excitation spreads through glutamatergic neurotransmission; and that rhythmic spontaneous burst events are disinhibited by morphine exposure.

**Fig. 6.**
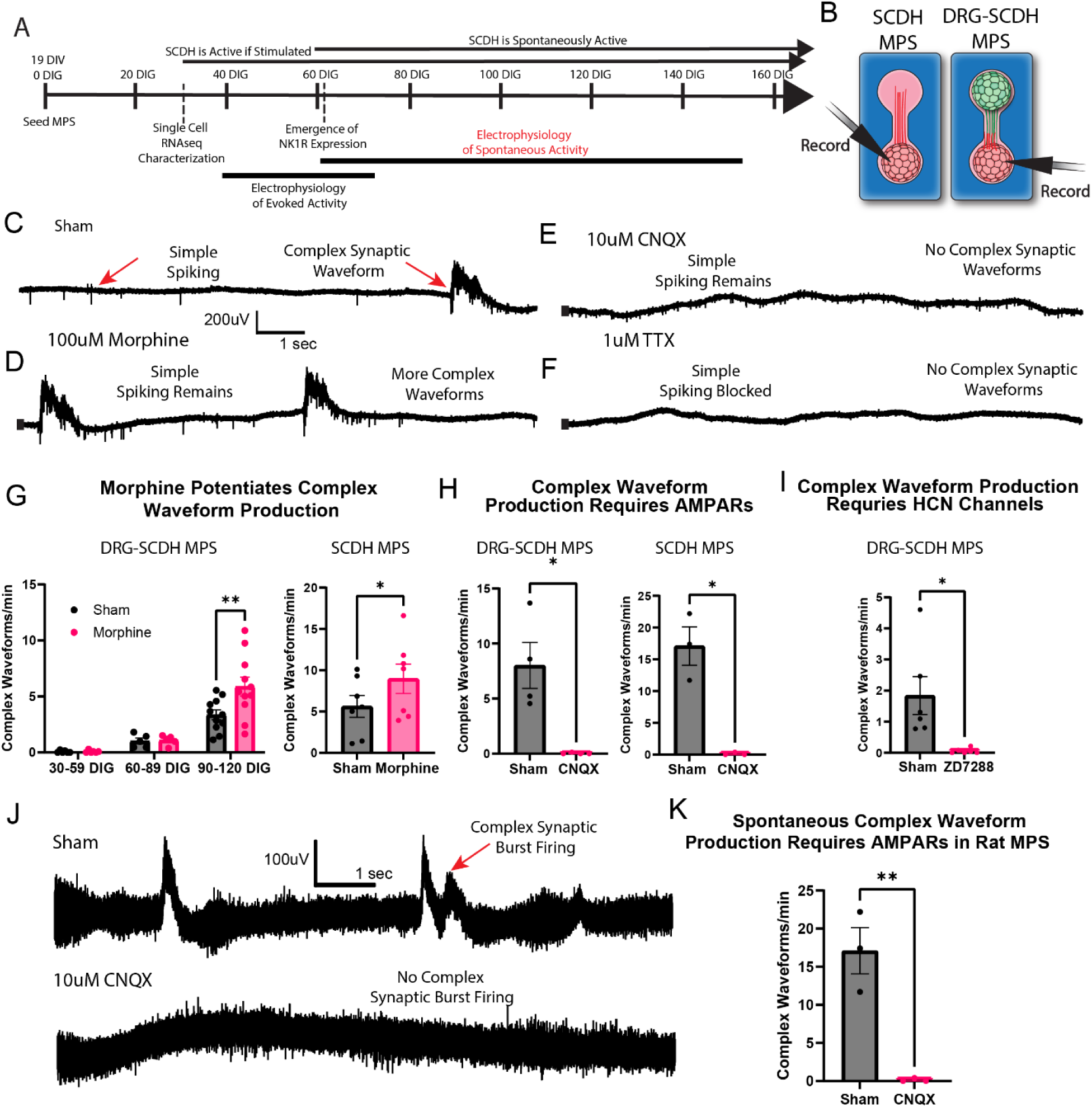
Spontaneous SCDH activity is disinhibited by morphine. **(A)** Spontaneous activity in the SCDH was evaluated between 70 and 120 DIG when this type of activity is consistently present. **(B)** SCDH field potentials were passively recorded in SCDH MPSs and DRG-SCDH MPSs in absence of any exogenous stimulation **(C)** Both simple spiking and fully spontaneous complex synaptic waveform production can be observed in mature nerve cultures in complete absence of any exogenous electrical stimulation. **(D)** The frequency of spontaneous waveforms increases after application of 100µm morphine. **(E)** Complex synaptic waveform production is abolished after inhibition of AMPAR-dependent glutamatergic synaptic transmission with CNQX. **(F)** All activity is abolished after application of the sodium channel blocker tetrodotoxin (TTX). **(G)** Application of morphine increases the frequency of complex waveform production inhibition whether SCDH tissue is innervated by nociceptor tissue or cultured in isolation. **(H)** Complex waveform production is abolished by AMPAR inhibition whether SCDH tissue is innervated by nociceptor tissue or cultured in isolation. **(I)** In innervated SCDH cultures, complex waveform production is abolished after incubation with the HCN channel blocker, ZD7288. **(J)** Similar complex synaptic waveforms are observed in an innervated SCDH microphysiological system constructed with embryonic rat DRG and SCDH tissue, which are **(K)** similarly abolished by AMPAR inhibition. **p<0.01 per a 2-way mixed-model ANOVA with LSD post-hoc test (E). *p<0.05, **p<0.01 per a paired t test (F,G,I).

## DISCUSSION

### Novel SCDH and human DRG-SCDH MPS derivation methodology

Starting with our previously published deterministic HOX patterning protocol to generate cervicothoracic spinal progenitors (*36*), we have demonstrated a novel method to generate transcriptionally and functionally appropriate hPSC-derived SCDH-IN cultures. These 3D neural cultures include critical late-born NK1R+ neurons that play a crucial role in pain and itch signaling and are potential treatment targets (*6, 17*). Unlike a prior hPSC-derived, 4-organoid, assembloid model containing DRG sensory neuron-NK1R+ SCDH neuron synapse (*29*), our DRG-SCDH connectoids can be derived in a scalable ‘off-the-shelf’ manner from cryopreserved cell banks (Fig. 2A). Moreover, combined with a recently commercialized MEA well plate (*52, 53*), our methodology could be directly translated as hPSC-derived novel alternative method (NAM) to model pain and itch chronicity and screen for potential therapeutics at scale.

In our SCDH spheroids, late-born neuronal types emerge without specific exposure to additional exogenous differentiation factors but do require extended culture periods (Fig. 3H,I). MPS are ideally suited for long-term culture of neuronal tissues, and we show that the 3D spheroid format permits the maintenance of a Sox2+/Pax6+ neural progenitor pool that enables derivation of late-born neuronal subtypes (Fig. 2D). This mirrors neural organoid technology in which certain cell types only emerge after many months in organoid culture (*54, 55*). Future investigation of the microenvironmental signals that govern the progenitors’ early-to-late born transition is warranted and experimentally tractable using our SCDH MPS.

### Expression of pain and itch-related genes in DRG-SCDH MPS

A recent unbiased transcriptomic analysis identified at least 15 distinct types of glutamatergic and 15 distinct types of GABAergic neurons in the mouse dorsal spinal cord (*56*). We demonstrate that prolonged MPS culture produces a greater diversity of these SCDH-INs than analogous 2D derivation/culture (Fig. 3C). We did not observe extensive expression of neuromodulator receptor genes at 35 DIG but we confirmed that neurons expressing the substance P receptor (NK1R) emerge at later timepoints that coincide with the emergence of spontaneous SCDH circuit activity (Fig. 4H,I and Fig. 5). We also demonstrate that MPS-derived interneuron populations express a wider array of dorsal interneuron identity markers including *FOXD3, PAX8, and BHLHE22* none of which were expressed to any significant extent following rapid 2D differentiation alone (Fig. 3C). Of note, the *BHLHE22* gene produces a transcription factor (also known as Bhlh5) that is transiently expressed during embryonic and early postnatal development but is required to initiate a developmental trajectory that produces a specific population of dorsal horn inhibitory neurons, i.e., the B5-I interneurons. These interneurons are characterized by expression of dynorphin or nNos, and the loss of these neurons through genetic ablation of the Bhlhb5 transcription factor induces a chronically pruritic phenotype in rodents (*13, 15*).

Gate control theory has long postulated that inhibitory (GABAergic) dorsal interneurons provide tonic inhibition to dorsal spinal circuitry that modulate ascending somatosensory information before it is projected to the brain for conscious perception (*10*). Endogenous release of dynorphins is thought to locally regulate the excitability of dorsal spinal circuits to modulate the perception of pain (*13*) but these inhibitory interneurons are also sensitive to exogenous opioids. In both the rodent dorsal spinal cord and our MPS’s SCDH, opioid receptors are expressed evenly by glutamatergic excitatory interneurons and GABAergic inhibitory interneurons, with the mu opioid receptor being most prevalent, followed by the kappa opioid and minimal expression of the delta opioid receptor (Fig. 3G) (*57, 58*). Morphine-dependent depression of GABAergic cell activity, and disinhibition of downstream excitatory cells types in the dorsal spinal cord, has been described in mouse (*12, 59*) and rat (*60*). Here, we extend these results to our human DRG-SCDH MPS (Fig. 5,6). Furthermore, DREADD silencing of GABAergic spinal cord neurons induced nocifensive behaviors and occluded subsequent effects of morphine (*32*), indicating that morphine’s actions in the spinal cord are mediated via GABAergic neurons.

In addition to its effect on nociception, chronic opioid exposure often induces chronic pruritus and even paradoxical pain hypersensitization (*61*). The pruritic effects of opioids are especially pronounced when the analgesic is delivered neuraxially (*21*). Inhibition of tonically active GABAergic neurons, and resulting disinhibition of SCDH circuits, is a plausible mechanism of both opioid-induced pruritus and paradoxical pain hypersensitization.

### Relevance of the SCDH-DRG MPS’s electrophysiological behavior

We show that morphine application to our microphysiological model of innervated spinal cord has a net excitatory effect on both evoked and spontaneous bioelectric activity in the SCDH spheroid. To our knowledge, we are the first to show the sensitivity of these spontaneous patterns of activity to morphine in a hPSC-derived model. The disinhibitory effects of morphine were specific to SCDH spheroid and strikingly similar to physiological effects described in an analogous embryonic rat model we described previously (*40*).

Synchronous spontaneous burst firing patterns have been described in rodent dorsal spinal cord slice preps and were shown to be altered under conditions of chronic pain and during exposure to analgesics (*50, 51, 62, 63*). These patterns are believed to be ultimately initiated by spontaneously-firing, pattern-generating cells that require hyperpolarization-induced cation currents, which can be inhibited by ZD7288 exposure, and continue independent of synaptic transmission at the single cell level (*51*). These pattern generators then drive rhythmic activity in a set of interconnected neural circuits composed of a wide variety of glutamatergic and GABAergic neurons that differ in regularity of rhythmic firing, the length of burst firing periods, number of spikes fired with each burst, consistency of burst firing frequencies, sensitivity to synaptic inhibition, and sensitivity to distinct sensory modalities (*51, 56, 64, 65*). This two compartment SCDH-DRG system shows that the fundamental properties of spontaneous, rhythmic firing of complex synaptic circuits are generated within SCDH spheroids, independent of any ascending or descending inputs provided *in vivo* or in previously published sensory assembloids that included cortical and thalamic compartments (Kim et al.) (Fig. 6). Moreover, the rhythmic firing is sensitive to ZD7288 exposure indicating the potential presence of HCN-expressing pattern-generating cells (Fig. 6I).

Collectively, our cellular phenotype and electrophysiology results support the utility of using our hiPSC-derived DRG-SCDH connectoids to study opioid-dependent changes in excitability of pain and itch-related spinal circuits. Differentiation of hiPSC lines could be used to study sex- specific and patient-specific differences in opioid responsiveness. Inclusion of immunocytes, particularly microglia, could potentially produce a sex and patient-specific model of opioid- dependent effects on neuro-immune interactions similar to our recent description in a peripheral nerve MPS (*66*). We have also recently demonstrated the ability to fabricate this innervated spinal cord MPS on a built-in microelectrode array (*52*) which will significantly increase experimental throughput and the breadth of questions that can be answered with this model.

## MATERIALS AND METHODS

### Study Design

Human SCDH and peripheral sensory neurons were derived in large numbers, aliquoted, and cryopreserved. MPSs were constructed in batch-wise fashion from these aliquots. In total, the data presented here was collected and compiled across six batches of MPS constructs all generated from aliquots of the same original population of differentiated peripheral sensory and SCDH neurons. Cells from two batches of MPS nerve constructs were pooled to generate the presented single-cell RNAseq, qPCR, and neurodevelopmental immunofluorescence data dataset. Recordings from four distinct batches of MPS nerve constructs were compiled to generate the electrophysiological data sets presented here. Both evoked (40-70 DIG) and spontaneous (70+ DIG) electrophysiological experiments were performed over time within each of the four batches. Immunofluorescent analysis of gross MPS tissue structure (Fig. 2C) was performed on a single batch of constructs.

### Human Stem Cell-Derived Dorsal Horn Differentiation

Experiments were conducted using the iPS(IMR90)-4 (WiCell) hiPSC line in feeder-free conditions maintained and differentiated (*36*) as previously described. To initiate differentiation, confluent hPSCs (70-90%) were singularized with Accutase (Invitrogen) and replated onto 35 mm Matrigel^®^-coated plates at a density of 1.5 x 10^5^ cells/cm^2^ in E8 medium with 10 μM ROCK inhibitor (Y27632, Tocris). The media was replaced with Essential 6 (E6) on the following day (Day 0). On Day 1, E6 media was supplemented with 200 ng/mL FGF8b (Peprotech). On Day 2, *HOX* propagation was initiated by sub-culturing at a 2:3 ratio and addition of neuromesodermal progenitor (NMP) media (E6 media with 200 ng/mL FGF8b and 3 μM CHIR99021 (Tocris)). Cells were washed once with PBS, incubated in Accutase for 1.5-2 mins, and removed from the surface by gentle pipetting. After centrifugation, cells were gently resuspended in NMP Medium containing 10 μM Y27632 and seeded on 35 mm Matrigel-coated plates. NMP Media was replenished on Day 4. Cells were sub-cultured again at a 2:3 ratio on Day 5. On Day 7, cultures were switched to spinal progenitor (pCNSP) media (E6 with 1 µM retinoic acid (RA; Sigma), 10 µM SB-431542 (Abcam), and 100 nM µL LDN-193189 (Stemgent)). On Day 8, cells were singularized and replated at 5 x 10^5^ cells/cm^2^ in pCNSP media containing 10 μM Y27632 for an additional 2 days. From Day 9-13, cultures were differentiated to dorsal horn progenitors in E6 media with 1 μM RA, then rapidly differentiated to neurons from Day 13-19 in maturation media (E6 with 1x N2 Supplement (Thermofisher), 1x B27 Supplement (Thermofisher), 1 µM cAMP (Sigma), 10 ng/mL GDNF, 10 ng/mL BDNF, 10 ng/mL NT-3 (Peprotech), and 10 µM DAPT (Tocris)). Dorsal horn neurons were cryopreserved on day 19 prior to downstream applications and analyses as previously described (*36*).

### Immunocytochemistry of Differentiating Neurons

For immunocytochemical analyses of frozen banks, cells were thawed and plated onto chamber slides in maturation media containing 10 μM Y27632 for 3 days. Cultures were then fixed in 4% paraformaldehyde for 10 mins, washed three times in PBS, and blocked in tris-buffered saline (TBS) containing 0.3% Triton-X 100 and 5% normal donkey serum (TBSDT) for one hour. The cells were incubated in primary antibodies (Table S1) diluted in TBSDT overnight at 4°C. After three 15 min washes in TBS containing 0.3% Triton-X 100, the cells were incubated with Alexa Fluor secondary antibodies (Invitrogen) at a 1:500 dilution in TBSDT for one hour at room temperature. Cells were washed twice in TBS for 15 mins each, counterstained with 300 nM DAPI for 10 mins, and washed once more in TBS prior to mounting with ProLong Gold Antifade (Life Technologies). Images were acquired using a Nikon A1R confocal microscope with Nikon NIS-Elements software and analyzed with NIS-Elements and ImageJ.

### Cell Aggregation

Cryopreserved human iPSC-derived peripheral sensory neurons were purchased from Anatomic Inc. (RealDRG) (Minneapolis, MN) (*28, 33*) and SCDH-INs were derived as described here (Fig. 1C) and previously (*36*). Cells were quickly thawed, diluted dropwise with Neurobasal media, centrifuged 300 x g for 4 min, and resuspended in human coculture media (neurobasal media containing 2% B27 supplement, 1% N2 supplement, 1% anti-anti, 1% glutamax, 10 ng/mL BDNF, 10 ng/mL GDNF, 20 ng/mL NGF, 10 ng/mL NT3, 1 µM dibutyrl-cAMP). RealDRG neurons and SCDH-INs were plated separately on Matrigel^®^ (Corning, NY, USA) coated T25 or 6-well culture plates flasks at a density of 60,000 cells/cm^2^ and allowed to adhere and recover overnight in human coculture media.

The following day, adherent human iPSC-derived SCDH-INs and RealDRG neurons were gently washed with pre-warmed media to removed dead cells and lifted by incubation in AccuMax™ or Accutase for 15 min at 37℃. Human coculture media was added to quench the Accutase reaction and the cells were gently triturated to create a single cells suspension. Cells were then pelleted for 4 minutes at 300 x g, the pellet was washed with 1.5mL culture media, and the cells were centrifuged again for 4 minutes at 300 x g. The supernatant was removed and cells were gently resuspended in fresh human coculture media and counted on a hemacytometer or on a Countess^TM^ automated cell counter (ThermoFisher). The cell suspension was diluted to the appropriate concentration and seeded into ultra-low attachment round-bottom 96-well plates at a density of 25k cells/well in 200 µL of human coculture media. The plates were then centrifuged for 5 minutes at 500 x g and kept at 37°C for 48 hours to allow for spheroid formation.

### Hydrogel Micropatterning

Fabrication and validation of 3D dual-hydrogel nerve cultures have been extensively described (*40*). Growth-restrictive outer-gels were first fabricated on polyethylene terephthalate Transwell^®^ membranes with 0.4 µm pore diameter (Corning Inc., NY) by photo-crosslinking a solution of 10% w/v polyethylene glycol dimethacrylate (PEG, Polysciences, PA), 1.1 mM lithium phenyl- 2,4,6-trimethylbenzoylphosphinate (LAP; Allevi, PA), and 0.0001% w/v TEMPO (Millipore- Sigma, MO) in phosphate-buffered saline (PBS, pH 7.4) with ultraviolet (UV) light patterned to create a long, thin inner void, with a bulb on one or both ends for spheroid placement, surrounded by a growth-restrictive PEG mold. Final bulbs measure 1 mm in diameter, are spaced 2mm from center of bulb to center of bulb, and are connected by a growth channel that is 0.4 mm wide.

### MPS Culture Maturation

To create DRG MPSs, a single DRG spheroid was placed in one bulb of the mold while the other was left empty. To create SCDH MPSs, a single SCDH spheroid was placed in the bulb of the mold while the other was left empty. To create innervated SCDH MPSs, a DRG spheroid was placed in the bulb at one end, and an SCDH spheroid was placed in the bulb at the other end of the same mold. All inner voids were then filled with a growth-permissive inner-gel solution composed of Matrigel^®^(Corning, Corning, NY) diluted 1:1 with human coculture media and incubated for 15 minutes at 37°C to allow for Matrigel^®^ gelation. Culture media was then added underneath Transwell^®^ membranes and cultures were returned to the incubator for maturation. Assembled dual-hydrogel MPS were matured for 30-225 days in gel (DIG) before evaluation, with media changes on Mondays, Wednesdays, and Fridays.

### Single Cell Dissociation of MPS

Cell spheroids were manually removed from the MPS devices and singularized for scRNAseq by dissociation with Papain (Worthington). Using sterilized forceps, cell spheroids were removed from the PEG hydrogel mold, transferred to a conical tube containing fresh media, and allowed to settle for 10 minutes. The spheroids were then resuspended in Papain solution containing DNase and incubated for 1.5 hours at 37° C on an orbital shaker. Cells were then triturated vigorously, centrifuged for 5 minutes at 300 x g, and quenched in ovomucoid solution containing DNase. Quenched cells were centrifuged, gently resuspended in PBS containing 0.2% BSA and 10 μM Y27632, then passed through a 40 µm cell strainer (Mitenyi Biotec) to remove debris. Cells were quantified and diluted to 700 cells/mL for sequencing.

### Single Cell RNA-Sequencing of MPS

9000 cells were targeted for capture from each sample. Transcriptomic profiling was performed using the Chromium Single Cell Gene Expression system (10X Genomics), according to the manufacturer’s recommendations using the Single Cell 30 Reagent v2/v3 kits (10X Genomics). Post-GEM-RT and post-cDNA amplification cleanup were performed using Dynabeads MyOne silane beads (Thermofisher) and SPRIselect (Beckman Coulter) kits respectively. Successful library preparation was confirmed using an Agilent Bioanalyzer (High Sensitivity DNA kit) and Qubit Fluorometer (High Sensitivity dsDNA kit). Experiment data was demultiplexed using the Cell Ranger Single Cell Software Suite, mkfastq command wrapped around Illumina’s bcl2fastq. The MiSeq balancing run was quality controlled using calculations based on UMI-tools (Smith, Heger and Sudbery, 2017). Samples libraries were balanced for the number of estimated reads per cell and run on an Illumina NovaSeq 6000 system. Cell Ranger software was then used to perform demultiplexing, alignment, filtering, barcode counting, UMI counting, and gene expression estimation for each sample according to the 10x Genomics documentation (https://support.10xgenomics.com/single-cell-gene-expression/software/pipelines/latest/what-is-cell-ranger). The gene expression estimates from each sample were then aggregated using Cell Ranger (cellranger aggr) and processed through the 10X Genomics Loupe Browser (v.6.3.0) for UMI filtering, cell clustering, and gene expression analyses. Genes that appeared in fewer than 5 cells and cells with fewer than 5000 UMIs were filtered from the dataset. Further analysis and visualization was performed using Loupe Browser (10X Genomics) and the Seurat toolkit in R. scRNA-seq data are available at GSE303698 and GSE186697

### Electrically-Evoked Neurophysiological Evaluation

Cultures were matured for 40-75 days prior to analysis of electrically-evoked circuit physiology. Mature cultures were removed from the incubator and placed in the electrophysiological recording apparatus. Under a microscope, the tip of a platinum wire (A-M Systems, Sequim, WA) inside a pulled-glass field recording electrode with 1 MΩ resistance was inserted into the recording site. Field potentials were recorded from the DRG or SCDH spheroid region in MPS cultures and the SCDH spheroid region of innervated nerve cultures. Next, a concentric bipolar stimulating electrode (FHC Inc., Bowdoin, ME) was inserted into the desired stimulation site (the nerve tissue 2 mm distal from the neural spheroid when recording from isolated DRG or SCDH tissue and the DRG spheroid of innervated SCDH cultures (Figs. 4A, 5B, 6B). Cultures were allowed to equilibrate in sham recording solution (ACSF or 0.1% DMSO in ACSF) for 30 minutes at room temperature prior to data collection, refreshing the ACSF every ten minutes. Ten replicate 200- µsec, bipolar, square wave 40V amplitude, spaced 20 seconds apart were then sequentially delivered to the stimulation site using LabChart Software (AD Instruments, Colorado Springs, CO). The resulting field potentials recorded at the recording site were amplified with a Model 3000 AC/DC Differential Amplifier (A-M systems, Sequim, WA) set at 100X gain and 0.1 Hz high-pass and 3 kHz low-pass filtering, and electrical interference was removed with a Hum Bug Noise Eliminator (Quest Scientific, North Vancouver, Canada). Traces were digitized with a PowerLab analog-to-digital converter (AD Instruments, Sidney, Australia) and recorded in LabChart. The sham recording solution was then replaced with the appropriate treatment solution. Cultures were incubated in treatment solution for ten minutes. Then, the treatment solution was refreshed, and the stimulation protocol was repeated.

The ten traces obtained under sham conditions were averaged into a single trace. The amplitude of each of the individual waveform components were calculated. As shown in Fig. 4B, D, and F and consistent with previous work in an analogous embryonic rat-based MPS (*40*), the N1 peak was determined to be the first sharp, negative-going peak observed after stimulation. The P1-early amplitude was determined to be the first sharp, positive-going peak observed after N1. The N2 peak was the largest amplitude negative-going peak between N1 and P1-early. The P1-late was determined to be the maximum value of the first broad, positive-going peak following P1-early and any additional broad, positive-going peaks following P1-late were counted as P2, P3, etc. Additional broad, positive-going peaks were observed to be disconnected from the electrically-evoked response in the SCDH and were therefore considered pseudo-spontaneous waveforms. The change in amplitude of N1, N2, P1-early, and P1-late between sham and treated conditions was calculated and statistical significance was determined with a one sample t test with a theoretical mean of zero indicating no change. The fold change in the number of semi-spontaneous waveforms between sham and treated conditions was calculated and statistical significance was determined with a one sample t test with a theoretical mean of 1 indicating no change.

### Spontaneous Neurophysiological Evaluation

When analyzing spontaneous field potential production, mature cultures were removed from the incubator and placed in the electrophysiological recording apparatus and bathed in sham recording solution or treatment solution. Under a microscope, the tip of a platinum wire inside a pulled-glass field recording electrode with 1 MΩ resistance was inserted into the recording site. Again, field potentials were recorded from the DRG or SCDH spheroid region in isolated nerve cultures and the SCDH spheroid region of innervated nerve cultures. Next, a concentric bipolar stimulating electrode (FHC Inc., Bowdoin, ME) was inserted into the desired stimulation site but was not used. Field potentials were passively recorded at the recording site for one hour while refreshing the recording solution every ten minutes.

The HCN channel inhibitor, ZD7288, binds to an intracellular bindings site that requires cellular uptake to be effective. This requires the drug to be incubated with neurons for one hour prior to observation of any effect (*50, 51*). Preliminary experiments determined that this mechanism was applicable to our MPS, therefore ZD7288 was applied to cultures in the incubator for one hour prior to the recording session.

The number of distinct complex waveforms produced across the hour-long recording session were counted. Counts from treated cultures were paired with the sham control recorded on the same day and statistical significance was determined with a 2-way mixed model ANOVA (Treatment x Time, repeated measures on Treatment) or a paired t-test where appropriate.

### Treatment Solutions

1 mM or 100 µM morphine sulfate salt, pentahydrate (Millipore-Sigma) dissolved directly in ACSF. 10 µM 6-cyano-7-nitroquinoxaline-2,3-dione (CNQX, Millipore-Sigma) dissolved in 0.1% DMSO. 100 µM bicuculline (Millipore-Sigma) dissolved in 0.1% DMSO. 1 µM tetrodotoxin (TTX, Abcam) dissolved in 0.1% DMSO. 100 µM ZD7288 (Millipore-Sigma) dissolved in 0.1% DMSO.

### qPCR

Total RNA was harvested in Trizol (Invitrogen) and isolated following the manufacture instructions of the Qiagen RNeasy Lipid Tissue Mini Kit. cDNA was generated using the SuperScript IV First-Strand Synthesis System (Invitrogen) according to the manufacturer’s instructions. TaqMan Gene Expression Assays (Table S2) and TaqMan Gene Expression Master Mix (Applied Biosystems) were used on a Bio-Rad CFX96 thermocycler with the following protocol: 50°C for 2 min; 95°C for 10 min; 40 cycles of 95°C for 15 s and 60°C for 1 min. Gene-expression was normalized to RPS18, and comparative fold-change analysis was performed using ΔCt.

### Whole-Culture Immunofluorescent Staining

Cultures were washed 6×10 minutes in PBS and fixed overnight at room temperature in excess of 2% paraformaldehyde (Electron Microscopy Sciences, Hatfield, PA, USA) with no agitation. Cultures were washed for 3×10 minutes, then blocked and permeabilized for 1 hour in 5% goat or donkey serum and 0.3% Triton-X 100-100 in PBS. Primary antibodies (Table S1) were diluted in 5% goat or donkey serum and 0.3% Triton-X 100-100 in PBS, applied to cultures, and incubated overnight at 4°C. Excess primary antibody was removed using 3×10 minute washes in PBS. Secondary antibodies were then diluted in 5% goat serum in PBS, applied to cultures, and incubated overnight at 4°C. Cultures were incubated in 2.5 µg/mL DAPI solution for 30 minutes at room temperature before being washed 3×10 minutes in PBS to remove excess secondary antibody. Alternatively, DAPI incubation was omitted, and cultures were washed 3×10 minutes in PBS directly after secondary incubation. Cultures were then manually transferred to microscope slides for mounting. Cultures were submerged in ProLong Glass Antifade Mountant with or without NucBlue (ThermoFisher), covered with a coverslip, and cured overnight at room temperature.

### 3D Confocal Microscopy

Confocal microscopy was performed on an Olympus IX 83 inverted confocal laser scanning microscope operated through the Olympus FluoView image acquisition software or on a Nikon AR1 confocal operated using Nikon NIS software. The appropriate 10x or 20x field of view was selected, and a z-stack containing the full depth of the tissue in that region was collected using optimal lateral and axial Nyquist criteria. The brightest plane of focus was identified, and acquisition settings were adjusted to minimize saturation. A 3D scanning region encompassing the entire stain construct was defined, images were acquired, and z-stacks were stitched together into a single 3D image. Singles planes of focus or maximum intensity projections were prepared as necessary.

### Fabrication of Embryonic Rat-Based MPS

An analogous rat MPS has been described extensively (*40*). All animal handling and tissue harvesting procedures were performed according to guidelines set by the U.S. NIH and approved in advance by the Institutional Animal Care and Use Committee (IACUC) at Tulane University. All DRG and SCDH tissue from a litter of rat fetuses at embryonic day 15 was harvested, pooled, and digested in 0.25% trypsin in phosphate-buffered saline (PBS), pH 7.4, at 37°C for 15 minutes. Tissue was then pelleted at 500×g for five minutes, trypsin was removed, and tissue was gently resuspended in rat coculture media (Neurobasal Medium supplemented with 2% v/v B27 supplement, 1% v/v N2 supplement, 1% v/v GlutaMAX, 20 ng/mL nerve growth factor 2.5S native mouse protein, 10 ng/mL recombinant human/murine/rat brain derived neurotrophic factor (PeproTech, Cranbury, NJ, USA), 10 ng/mL recombinant human glial cell derived neurotrophic factor (PeproTech), and 1% v/v antibiotic/antimycotic solution (all from Thermo-Fisher Scientific, Waltham, MA unless otherwise noted)). Digested tissue was dissociated through trituration, and passed through a 40 µM nylon mesh filter. DRG and SCDH cells were then separately seeded in 96-well ultra-low attachment spheroid microplates at a concentration of 45,000 DRG cells per well and 60,000 SCDH cells per well in rat coculture media. Microplates were centrifuged at 500×g for five minutes and placed in the incubator to aggregate at 37°C and 5% CO_2_ for 48 hours. Resulting embryonic rat spheroids were seeded in hydrogels as described for hiPSC-derived nerve constructs.

### Statistics

Electrically-evoked electrophysiological analysis (Fig. 4C-G) could be performed with a repeated-measures design. The waveform was first evoked in vehicle (ACSF). Drug was applied and the waveform was evoked again. The amplitude of each waveform component was measured under vehicle and treated conditions and the change in amplitude was calculated. The average change in amplitude was then calculated for each waveform component for each drug. This average was then compared to a theoretical value of 0 (no change) with a one-sample t-test.

Spontaneous electrophysiological analysis was performed with a between-subjects design (Fig. 6G-I,K). Each construct was immediately bathed in either vehicle (ACSF) or drug dissolved in ACSF and passively recorded for at least an hour. Each drug-treated construct was paired with a vehicle-treated construct recorded on the same day. The frequency of spontaneous activity was calculated for each construct and averaged by treatment condition. Conditions were then compared with a mixed-model ANOVA (Fig. 6G, left) or a paired t-test (Fig. 6G, right, Fig. 6H,I,K).

## List of Supplementary Materials

Figure S1: scRNA-Seq Characterization of SCDH and DRG Neurons

Figure S2: Application of 1mM morphine depresses evoked bioelectric activity in innervated SCDH circuits, and SCDH and DRG spheriods cultured alone.

Figure S3: Application of 100μM morphine did not affect the frequency of spontaneous events that occurred in DRG-SCHD MPSs.

Table S1: Immunocytochemistry Reagents

Table S2: Taqman qPCR Gene Expression Assays

## Acknowledgements

We acknowledge the UW-Madison Biotechnology Center for all next-gen scRNA-sequencing including in this manuscript, and C. Birchmeier and T. Müller for gifted LBX1, TLX3, and LMX1B antibodies.

## Funding

National Institutes of Health grant UG3-UH3 TR0003150 (MJM, RA)

Louisiana Board of Regents Departmental Enhancement grant LEQSF[2018-23]-ENH-DE-15 (Tulane Brain Institute)

NSF Cell Manufacturing Technologies #1648035 (RA) NIH/NINDS F32 NS106740 (NI)

## Author contributions

Conceptualization: RSA, MJM

Methodology: KJP, FRS, NRI

Investigation: KJP, FRS, NRI

Visualization: KJP, FRS, NRI

Funding acquisition: RSA, MJM

Project administration: RSA, MJM

Writing – original draft: KJP, FRS, NRI

Writing – review & editing: KJP, FRS, NRI, AB, RSA, MJM

## Data and materials availability

All data needed to evaluate the conclusions in the paper are present in the paper and/or the Supplementary Materials. scRNA-seq data are available through the GEO repository (GSE303698 and GSE186697).

## References and Notes

1. H. Ikeda, B. Heinke, R. Ruscheweyh, J. Sandkuhler, Synaptic plasticity in spinal lamina I projection neurons that mediate hyperalgesia. Science 299, 1237–1240 (2003).

2. R. Wercberger, J. M. Braz, J. A. Weinrich, A. I. Basbaum, Pain and itch processing by subpopulations of molecularly diverse spinal and trigeminal projection neurons. Proc Natl Acad Sci U S A 118, (2021).

3. A. I. Nascimento, F. M. Mar, M. M. Sousa, The intriguing nature of dorsal root ganglion neurons: Linking structure with polarity and function. Prog Neurobiol 168, 86–103 (2018).

4. E. E. Benarroch, Dorsal horn circuitry: Complexity and implications for mechanisms of neuropathic pain. Neurology 86, 1060–1069 (2016).

5. S. E. Ross, J. Hachisuka, A. J. Todd, in Itch: Mechanisms and Treatment, E. Carstens, T. Akiyama, Eds. (Boca Raton (FL), 2014)

6. M. Gautam et al., Role of neurokinin type 1 receptor in nociception at the periphery and the spinal level in the rat. Spinal Cord 54, 172–182 (2016).

7. T. D. Sheahan, C. A. Warwick, L. G. Fanien, S. E. Ross, The Neurokinin-1 Receptor is Expressed with Gastrin-Releasing Peptide Receptor in Spinal Interneurons and Modulates Itch. J Neurosci 40, 8816–8830 (2020).

8. A. J. Todd, Neuronal circuitry for pain processing in the dorsal horn. Nat Rev Neurosci 11, 823–836 (2010).

9. H. C. Lai, R. P. Seal, J. E. Johnson, Making sense out of spinal cord somatosensory development. Development 143, 3434–3448 (2016).

10. R. Melzack, P. D. Wall, Pain mechanisms: A new theory: A gate control system modulates sensory input from the skin before it evokes pain perception and response. Science 150, 971–979 (1965).

11. C. Torsney, A. B. MacDermott, Disinhibition opens the gate to pathological pain signaling in superficial neurokinin 1 receptor-expressing neurons in rat spinal cord. J Neurosci 26, 1833–1843 (2006).

12. A. Francois et al., A Brainstem-Spinal Cord Inhibitory Circuit for Mechanical Pain Modulation by GABA and Enkephalins. Neuron 93, 822–839 e826 (2017).

13. A. P. Kardon et al., Dynorphin acts as a neuromodulator to inhibit itch in the dorsal horn of the spinal cord. Neuron 82, 573–586 (2014).

14. E. Foster et al., Targeted ablation, silencing, and activation establish glycinergic dorsal horn neurons as key components of a spinal gate for pain and itch. Neuron 85, 1289–1304 (2015).

15. S. E. Ross et al., Loss of inhibitory interneurons in the dorsal spinal cord and elevated itch in Bhlhb5 mutant mice. Neuron 65, 886–898 (2010).

16. J. M. Laird, C. Roza, C. De Felipe, S. P. Hunt, F. Cervero, Role of central and peripheral tachykinin NK1 receptors in capsaicin-induced pain and hyperalgesia in mice. Pain 90, 97–103 (2001).

17. P. Kleczkowska, K. Nowicka, M. Bujalska-Zadrozny, E. Hermans, Neurokinin-1 receptor-based bivalent drugs in pain management: The journey to nowhere? Pharmacol Ther 196, 44–58 (2019).

18. J. A. Trafton, C. Abbadie, K. Marek, A. I. Basbaum, Postsynaptic signaling via the [mu]-opioid receptor: responses of dorsal horn neurons to exogenous opioids and noxious stimulation. J Neurosci 20, 8578–8584 (2000).

19. CDC. (Centers for Disease Control, 2025).

20. P. Y. Ho, Y. C. Huang, Y. T. Lin, Systemic neurokinin-1 receptor antagonists mitigate chronic pruritus: A systematic review and meta-analysis. J Eur Acad Dermatol Venereol, (2025).

21. K. Kumar, S. I. Singh, Neuraxial opioid-induced pruritus: An update. J Anaesthesiol Clin Pharmacol 29, 303–307 (2013).

22. D. Tavares-Ferreira et al., Spatial transcriptomics of dorsal root ganglia identifies molecular signatures of human nociceptors. Science Translational Medicine 14, eabj8186.

23. R. Y. North et al., Electrophysiological and transcriptomic correlates of neuropathic pain in human dorsal root ganglion neurons. Brain 142, 1215–1226 (2019).

24. X. Li et al., Profiling spatiotemporal gene expression of the developing human spinal cord and implications for ependymoma origin. Nat Neurosci 26, 891–901 (2023).

25. T. Rayon, R. J. Maizels, C. Barrington, J. Briscoe, Single-cell transcriptome profiling of the human developing spinal cord reveals a conserved genetic programme with human-specific features. Development 148, (2021).

26. A. S. Yekkirala, D. P. Roberson, B. P. Bean, C. J. Woolf, Breaking barriers to novel analgesic drug development. Nat Rev Drug Discov 16, 545–564 (2017).

27. P. Roderer et al., Emergence of nociceptive functionality and opioid signaling in human induced pluripotent stem cell-derived sensory neurons. Pain 164, 1718–1733 (2023).

28. V. Truong et al., Transcriptomic analysis and high throughput functional characterization of human induced pluripotent stem cell derived sensory neurons. bioRxiv, (2024).

29. J. I. Kim et al., Human assembloid model of the ascending neural sensory pathway. Nature 642, 143–153 (2025).

30. E. Nguyen et al., Morphine acts on spinal dynorphin neurons to cause itch through disinhibition. Sci Transl Med 13, (2021).

31. Z. Wang et al., Central opioid receptors mediate morphine-induced itch and chronic itch via disinhibition. Brain 144, 665–681 (2021).

32. K. Koga et al., Chemogenetic silencing of GABAergic dorsal horn interneurons induces morphine-resistant spontaneous nocifensive behaviours. Sci Rep 7, 4739 (2017).

33. S. Jami et al., Pain-causing stinging nettle toxins target TMEM233 to modulate Na(V)1.7 function. Nat Commun 14, 2442 (2023).

34. P. J. Osseward, 2nd et al., Conserved genetic signatures parcellate cardinal spinal neuron classes into local and projection subsets. Science 372, 385–393 (2021).

35. N. E. Szabo et al., Hoxb8 intersection defines a role for Lmx1b in excitatory dorsal horn neuron development, spinofugal connectivity, and nociception. J Neurosci 35, 5233–5246 (2015).

36. N. R. Iyer et al., Modular derivation of diverse, regionally discrete human posterior CNS neurons enables discovery of transcriptomic patterns. Sci Adv 8, eabn7430 (2022).

37. K. Keino-Masu et al., Deleted in Colorectal Cancer (DCC) encodes a netrin receptor. Cell 87, 175–185 (1996).

38. C. Sabatier et al., The divergent Robo family protein rig-1/Robo3 is a negative regulator of slit responsiveness required for midline crossing by commissural axons. Cell 117, 157–169 (2004).

39. S. Gupta et al., Deriving Dorsal Spinal Sensory Interneurons from Human Pluripotent Stem Cells. Stem Cell Reports 10, 390–405 (2018).

40. K. J. Pollard, D. A. Bowser, W. A. Anderson, M. Meselhe, M. J. Moore, Morphine-sensitive synaptic transmission emerges in embryonic rat microphysiological model of lower afferent nociceptive signaling. Sci Adv 7, (2021).

41. J. C. Holstege et al., Loss of Hoxb8 alters spinal dorsal laminae and sensory responses in mice. Proc Natl Acad Sci U S A 105, 6338–6343 (2008).

42. S. Pachernegg, N. Strutz-Seebohm, M. Hollmann, GluN3 subunit-containing NMDA receptors: not just one-trick ponies. Trends Neurosci 35, 240–249 (2012).

43. M. A. Henson, A. C. Roberts, I. Perez-Otano, B. D. Philpot, Influence of the NR3A subunit on NMDA receptor functions. Prog Neurobiol 91, 23–37 (2010).

44. C. M. Niswender, P. J. Conn, Metabotropic glutamate receptors: physiology, pharmacology, and disease. Annu Rev Pharmacol Toxicol 50, 295–322 (2010).

45. L. E. Lorenzo et al., Enhancing neuronal chloride extrusion rescues alpha2/alpha3 GABA(A)-mediated analgesia in neuropathic pain. Nat Commun 11, 869 (2020).

46. M. J. Sykes et al., Neuron-specific responses to acetylcholine within the spinal dorsal horn circuits of rodent and primate. Neuropharmacology 198, 108755 (2021).

47. J. Li, M. L. Baccei, Cell type-dependent short-term plasticity and dopaminergic modulation of sensory synapses onto mouse superficial dorsal horn neurons. J Neurosci, (2025).

48. L. F. Borges, S. D. Iversen, Topography of choline acetyltransferase immunoreactive neurons and fibers in the rat spinal cord. Brain Res 362, 140–148 (1986).

49. A. Dahlstroem, K. Fuxe, Evidence for the Existence of Monoamine-Containing Neurons in the Central Nervous System. I. Demonstration of Monoamines in the Cell Bodies of Brain Stem Neurons. *Acta Physiol Scand Suppl*, SUPPL 232:231–255 (1964).

50. J. Lucas-Romero, I. Rivera-Arconada, J. A. Lopez-Garcia, Synchronous firing of dorsal horn neurons at the origin of dorsal root reflexes in naive and paw-inflamed mice. Front Cell Neurosci 16, 1004956 (2022).

51. J. Lucas-Romero, I. Rivera-Arconada, C. Roza, J. A. Lopez-Garcia, Origin and classification of spontaneous discharges in mouse superficial dorsal horn neurons. Sci Rep 8, 9735 (2018).

52. C. M. Didier et al., Fully Integrated 3D Microelectrode Arrays with Polydopamine-Mediated Silicon Dioxide Insulation for Electrophysiological Interrogation of a Novel 3D Human, Neural Microphysiological Construct. ACS Appl Mater Interfaces 15, 37157–37173 (2023).

53. C. Rountree et al., Human Peripheral Nerve-on-a-Chip on a Multiwell Microelectrode Array as a Scalable Preclinical Neurotoxicity Assay. bioRxiv, (2025).

54. M. A. Lancaster, J. A. Knoblich, Generation of cerebral organoids from human pluripotent stem cells. Nat Protoc 9, 2329–2340 (2014).

55. A. M. Pasca et al., Functional cortical neurons and astrocytes from human pluripotent stem cells in 3D culture. Nat Methods 12, 671–678 (2015).

56. M. Haring et al., Neuronal atlas of the dorsal horn defines its architecture and links sensory input to transcriptional cell types. Nat Neurosci 21, 869–880 (2018).

57. Z. Wang et al., Central opioid receptors mediate morphine-induced itch and chronic itch via disinhibition. Brain 144, 665–681 (2021).

58. Y. R. Kim, H. G. Shim, C.-E. Kim, S. J. Kim, The effect of µ-opioid receptor activation on GABAergic neurons in the spinal dorsal horn. The Korean Journal of Physiology & Pharmacology 22, (2018).

59. A. Kearns et al., Neuron Type-Dependent Synaptic Activity in the Spinal Dorsal Horn of Opioid-Induced Hyperalgesia Mouse Model. Front Synaptic Neurosci 13, 748929 (2021).

60. S. L. Jinks, E. Carstens, Superficial dorsal horn neurons identified by intracutaneous histamine: chemonociceptive responses and modulation by morphine. J Neurophysiol 84, 616–627 (2000).

61. A. Tinnermann, C. Sprenger, C. Büchel, Opioid analgesia alters corticospinal coupling along the descending pain system in healthy participants. eLife 11, (2022).

62. M. F. Bandres, J. Gomes, J. G. McPherson, Spontaneous Multimodal Neural Transmission Suggests That Adult Spinal Networks Maintain an Intrinsic State of Readiness to Execute Sensorimotor Behaviors. J Neurosci 41, 7978–7990 (2021).

63. J. Sandkuhler, A. A. Eblen-Zajjur, Identification and characterization of rhythmic nociceptive and non-nociceptive spinal dorsal horn neurons in the rat. Neuroscience 61, 991–1006 (1994).

64. A. Sathyamurthy et al., Massively Parallel Single Nucleus Transcriptional Profiling Defines Spinal Cord Neurons and Their Activity during Behavior. Cell Rep 22, 2216–2225 (2018).

65. S. J. Sullivan, A. D. Sdrulla, Excitatory and Inhibitory Neurons of the Spinal Cord Superficial Dorsal Horn Diverge in Their Somatosensory Responses and Plasticity in Vivo. J Neurosci 42, 1958–1973 (2022).

66. K. J. Pollard et al., Respiratory syncytial virus infects peripheral and spinal nerves and induces chemokine-mediated neuropathy. J Infect Dis, (2023).

