## Supplementary Information for "Human microphysiological model of dorsal root ganglion-spinal cord dorsal horn circuitry recapitulates opioid induced effects"

### Supplemental Figures

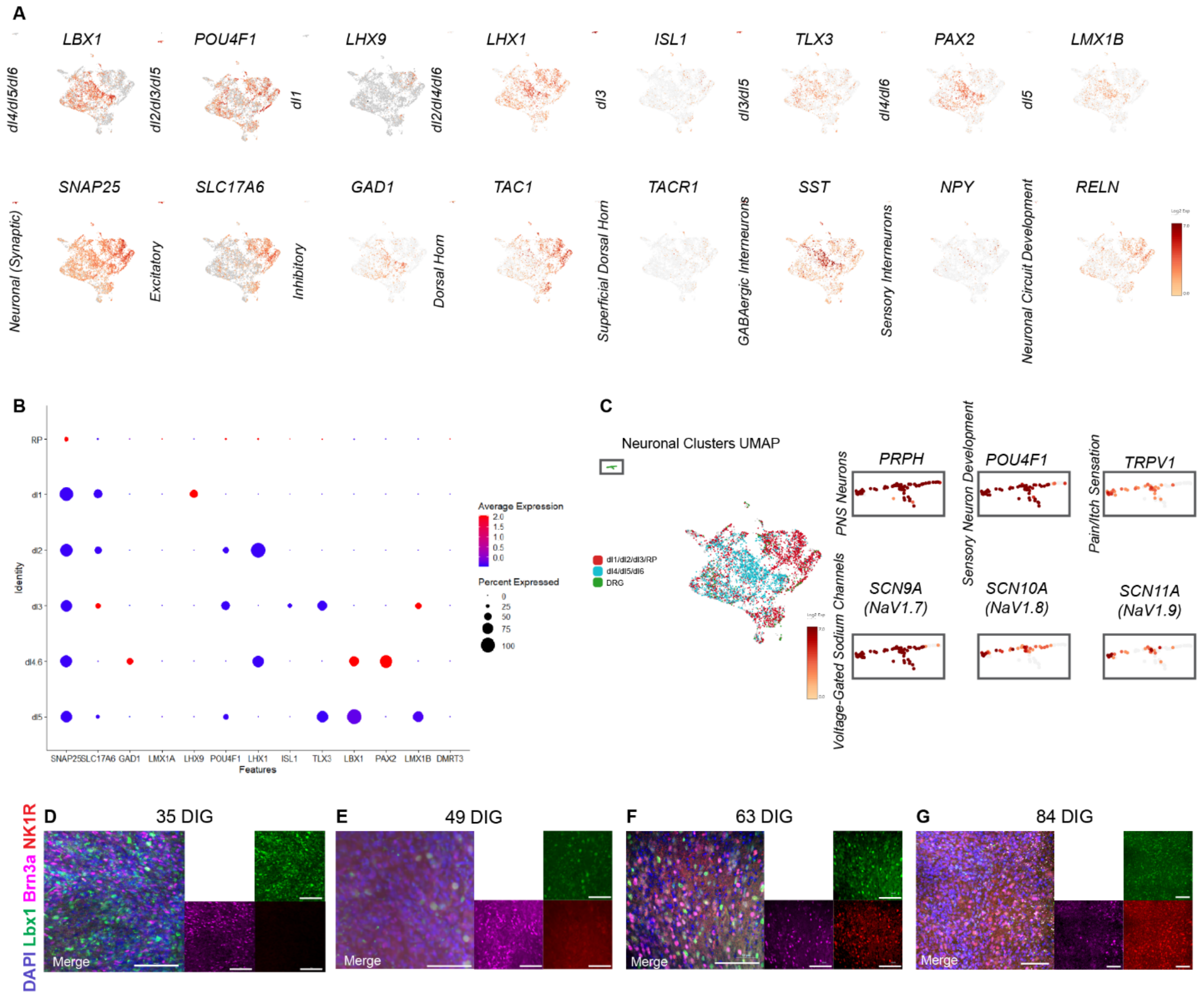


**Fig. S1. scRNA-Seq Characterization of SCDH and DRG Neurons: A)** Spinal cord neuronal subtype marker expression in 35 DIG MPS cultures highlighting the presence excitatory an inhibitory SCDH sub-populations. **B)** Dotplot indicating marker expression of 35 DIG SCDH-INs separated by subtype identity. Cells identified as dI4/6 have high expression of Lbx1, Pax2, and Lhx1 and cells identified as dI5 have high expression of Lbx1, Tlx3, Lmx1b, and Pou4f1. **C)** Nociceptor/pruriceptor expression in DRG cells from 35 DIG MPS cultures. Cells identified as DRG express markers for PNS neurons (PRPH) and sensory neuron development (POU4F1) as well as genes encoding ion channels characteristic of nociceptive and pruriceptive sensory neurons (TRPV1, NaV1.7, NaV1.8, NaV1.9). **D-G)** IHC of SCDH spheroids from the DRG-SCDH MPS at 35, 49, 63, and 84 DIG, respectively, indicating maintenance of SCDH-IN markers (Lbx1, Brn3a) throughout the culture time course. Widespread expression of NK1R can be seen in 63 and 84 DIG cultures as demonstrated in Figure 3.

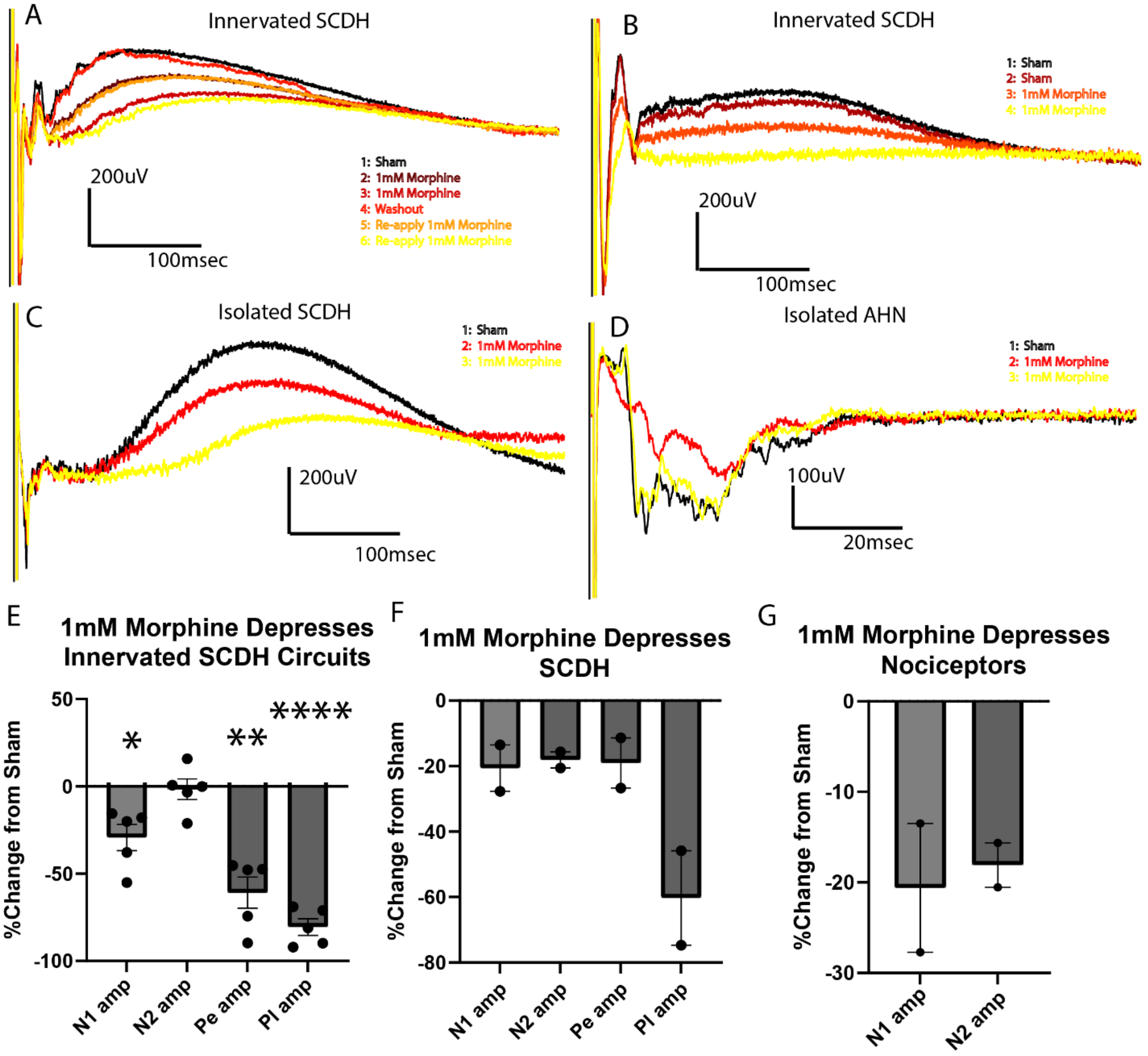


**Fig. S2**. **Application of 1mM morphine depresses evoked bioelectric activity in innervated SCDH circuits, SCDH and DRG spheroids cultured alone**. **(A,B, D)** The late positive-going field potentials recorded in SCDH after stimulation of innervating nociceptor tissue are significantly reduced following application of 1mM morphine, an effect that can be washed out and repeated. **(C,F)** Similar reduction in the amplitude of late, positive-going field potentials was observed when SCDH spheriod was cultured in insolation. **(E,G)** Late, positive-going field potentials are not observed following electrical stimulation of DRG spheroid, but application of 1mM morphine significantly reduced the early, negative-going electrical evoked field potentials suggesting that this concentration of morphine is also depressive to DRG neurons.

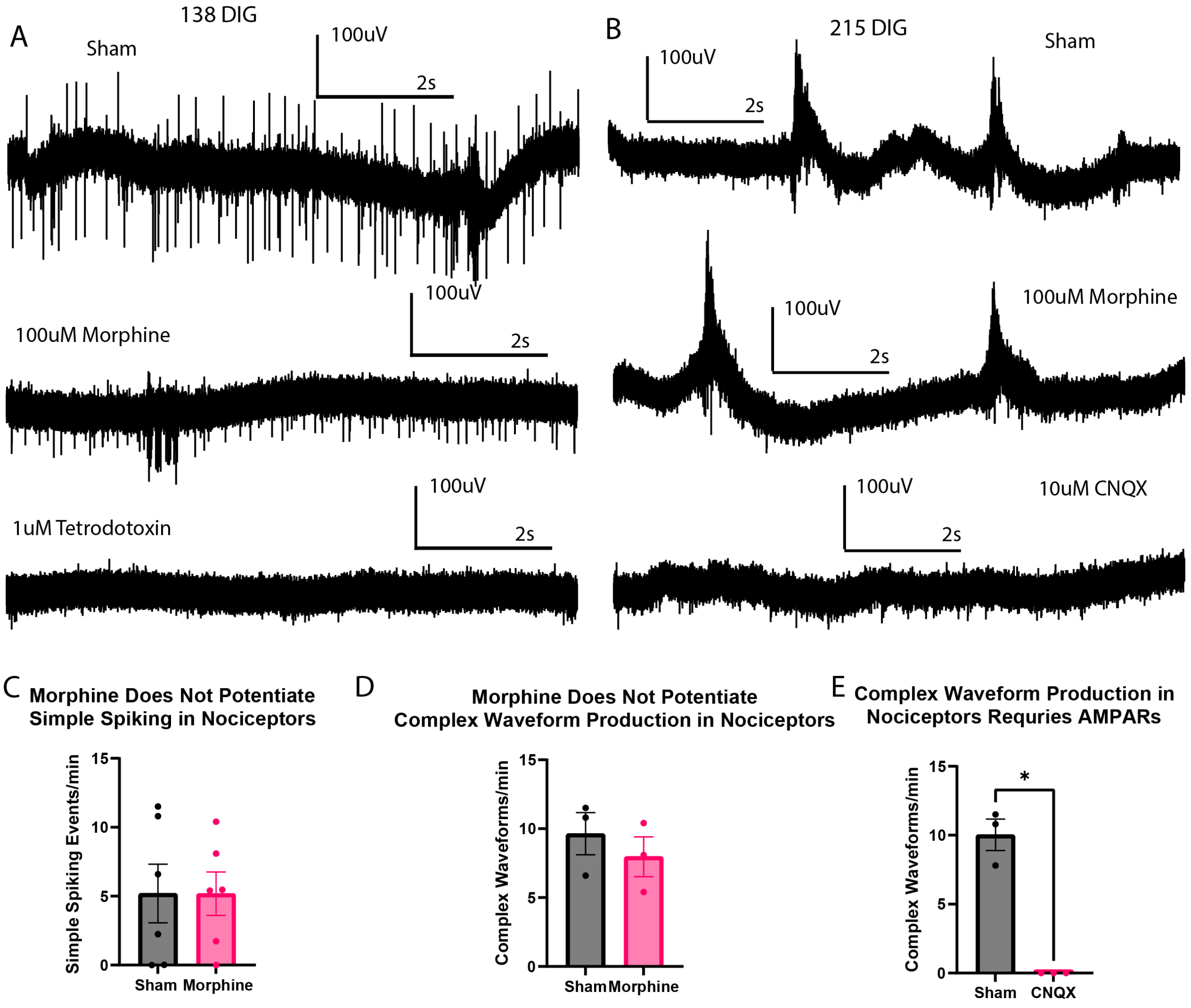


**Fig. S3**. **Application of 100μM morphine did not affect the frequency of spontaneous events that occurred in DRG-SCDH MPS. (A)** Simple spiking events were observed between 100 and 150 days in gel (DIG) while **(B)** complex waveform production was observed when cultures were allowed to mature for 215 DIG. **(C-D)** Application of 100μM morphine did not affect the frequency of simple spiking or complex waveform production. **(E)** Complex waveform production was abolished by application of the AMPA/Kainate receptor antagonist CNQX, indicating that they are driven by glutamatergic synaptic neurotransmission.

#### Supplemental Tables

| Species | Antigen | Dilution | Vendor | Catalog # |
|---|---|---|---|---|
| mouse | β-III-Tubulin | 1:1000 | Abcam | ab78078 |
| chicken | peripherin | 1:500 | ThermoFisher | PA110012 |
| chicken | GFAP | 1:1000 | Abcam | ab4674 |
| rabbit | Synapsin I | 1 µg/mL | Abcam | ab64581 |
| mouse | PSD95 | 1:500 | Neuromab | K28/43 |
| rabbit | Vgat | 1:100 | Novus Biologicals | NBP2-20857 |
| rabbit | Vglut2 | 1:250 | Abcam | ab216463 |
| rabbit | HCN1 | 1:100 | Abcam | ab229340 |
| mouse | Sox2 | 1:200 | Millipore | MAB4423 |
| mouse | NCadherin | 1:200 | BD Biosciences | 610920 |
| goat | Pax2 | 1:500 | R&D Systems | AF3364 |
| goat | Pax3 | 1:250 | R&D Systems | AF2457 |
| rabbit | Pax6 | 1:200 | Biolegend | 901301 |
| mouse | Pax7 | 1:20 | DSHB | PAX7 |
| guinea pig | Lbx1 | 1:200/*1:10,000 | Gift, C. Birchmeier and T. Mueller | RRID:AB_2532144 |
| mouse | Brn3a | 1:50 | SCBT | sc-8429 |
| goat | Isl1 | 1:500 | R&D Systems | AF1837 |
| rabbit | Tlx3 | 1:200/*1:10,000 | Gift, C. Birchmeier and T. Mueller | RRID:AB_2893157 |
| guinea pig | Lmx1b | 1:200/*1:10,000 | Gift, C. Birchmeier and T. Mueller | RRID:AB_2893158 |
| mouse | Lhx1/5 | 1:50 | DSHB | 4F2-c |
| goat | DCC | 1:200 | R&D Systems | AF844 |
| mouse | βIII-Tubulin | 1:1000 | Biolegend | 801202 |
| goat | Robo3 | 1:200 | R&D Systems | AF3076 |
| sheep | Onecut2 | 1:200 | R&D Systems | AF6294 |
| sheep | Zfhx3 | 1:200 | R&D Systems | AF7384 |
| rabbit | Nfia | 1:200 | Atlas Antibodies | HPA008884 |
| rabbit | NeuroD2 | 1:200 | Abcam | ab104430 |
| rabbit | Nk1R | 1:500 | Sigma | S8305 |

**Table S1. Primary antibody list:** This table describes each of the primary antibodies and antibody sources used in this study.

| Target | Vendor | Catalog# |
|---|---|---|
| RPS18 | Thermofisher | Hs01375212_g1 |
| Lmx1b | Thermofisher | Hs00158750_m1 |
| Lbx1 | Thermofisher | Hs00198080_m1 |
| NeuroD2 | Thermofisher | Hs00272055_s1 |
| Nfia | Thermofisher | Hs01374272_m1 |
| Zfhx3 | Thermofisher | Hs00199344_m1 |
| Onecut2 | Thermofisher | Hs00191477_m1 |
| TacR1 | Thermofisher | Hs00185530_m1 |

**Table S2. Taqman gene expression assay list:** This table describes each of the taqman gene expression assays and sources used in this study.
